# The Bouma law accounts for crowding in fifty observers

**DOI:** 10.1101/2021.04.12.439570

**Authors:** Jan W. Kurzawski, Augustin Burchell, Darshan Thapa, Jonathan Winawer, Najib J. Majaj, Denis G. Pelli

**Author notes:** Corresponding Author: Denis G. Pelli, Department of Psychology, 6 Washington Place, New York University, NY, 10003, USA.

## Abstract

*Crowding* is the failure to recognize an object due to surrounding clutter. Our visual crowding survey measured 13 crowding distances (or “critical spacings”) twice in each of 50 observers. The survey included three eccentricities (0, 5, and 10 deg), four cardinal meridians, two orientations (radial and tangential), and two fonts (Sloan and Pelli). The survey also tested foveal acuity, twice. Remarkably, fitting a two-parameter model, the well-known Bouma law — crowding distance grows linearly with eccentricity — explains 82% of the variance for all 13 × 50 measured log crowding distances, cross-validated. An enhanced Bouma law, with factors for meridian, crowding orientation, target kind, and observer, explains 94% of the variance, again cross-validated. These additional factors reveal several asymmetries, consistent with previous reports, which can be expressed as crowding- distance ratios: 0.62 horizontal:vertical, 0.79 lower:upper, 0.78 right:left, 0.55 tangential:radial, and 0.78 Sloan font:Pelli font. Across our observers, peripheral crowding is independent of foveal crowding and acuity. Evaluation of the Bouma factor *b* (the slope of the Bouma law) as a biomarker of visual health would be easier if there were a way to compare results across crowding studies that use different methods. We define a *standardized Bouma factor b’* that corrects for differences from Bouma’s 25 choice alternatives, 75% threshold criterion, and linearly symmetric flanker placement. For radial crowding on the right meridian, the standardized Bouma factor *b’* is 0.24 for this study, 0.35 for Bouma (1970), and 0.30 for the geometric mean across five representative modern studies, including this one, showing good agreement across labs, including Bouma’s. We found that guaranteeing fixation by gaze-contingent display halved the standard deviation across observers of the estimated log *b*. The reduction in standard deviation is explained by a “peeking” model in which the observer looked near an anticipated target location in 50% of *unmonitored*-fixation trials. Individual differences are robust, as evidenced by the much larger 0.08 SD of log *b* across observers than the 0.03 SD of test-retest within observers. Crowding’s ease of measurement enhances its promise as a biomarker for dyslexia and visual health.

## Introduction

Crowding is the failure to recognize an object due to surrounding clutter (Bouma, 1970, 1973; Pelli et al., 2004; Pelli & Tillman, 2008; Strasburger et al., 1991; Stuart & Burian, 1962). Crowding has been studied with several different tasks including letter identification (Bouma, 1970; Flom, Heath, et al., 1963; Strasburger et al., 1991), Landolt rings (Flom, Heath, et al., 1963; Flom, Weymouth, et al., 1963), Vernier acuity (Levi et al., 1985; Malania et al., 2007; Westheimer & Hauske, 1975), face recognition (Farzin et al., 2009; Louie et al., 2007; Martelli et al., 2005), and orientation discrimination (Andriessen & Bouma, 1976; Parkes et al., 2001; Toet & Levi, 1992; Westheimer et al., 1976). It is invariant with the size of target and flankers (Levi & Carney, 2009; Pelli et al., 2004; Pelli et al., 2007; Strasburger et al., 1991; Tripathy & Cavanagh, 2002). Crowding is usually measured by sandwiching the target between two similar flanking objects, or *flankers*, and is characterized by the *crowding distance* (or “critical spacing”), which is the center-to-center distance from target to flanker at which recognition attains a criterion level of performance. Crowding distance increases linearly with eccentricity (Bouma, 1970; Kooi et al., 1994; Levi & Carney, 2009; Pelli et al., 2004; Toet & Levi, 1992), and increases with target-flanker similarity (Andriessen & Bouma, 1976; Chastain, 1982; Kooi et al., 1994; Leat et al., 1999; Nazir, 1992; Pelli et al., 2004) as well as the number of distractors (Grainger et al., 2010; Strasburger et al., 1991).

Crowding also occurs for moving stimuli (Bex & Dakin, 2005; Bex et al., 2003). For review of the crowding literature see (Herzog et al., 2015; Levi, 2008; Pelli & Tillman, 2008; Strasburger, 2020; Strasburger et al., 2011; Whitney & Levi, 2011). Among the normally sighted, crowding was first reported in the periphery, and, after some debate, has now been convincingly demonstrated in the fovea (Atkinson et al., 1986; Coates et al., 2018; Flom, Heath, et al., 1963 1963; Liu & Arditi, 2000; Malania et al., 2007; Pelli et al., 2016; Siderov et al., 2013; Toet & Levi, 1992).

We are interested in relating psychophysical measures of crowding to brain physiology, especially cortical magnification measured by fMRI (functional Magnetic Resonance Imaging) in areas V1, V2, V3, and hV4. For this purpose, we tested crowding in 50 observers to characterize the statistics of crowding within and across individuals. The comparison with fMRI will be reported separately (Kurzawski et al., 2021). Here we report only the psychophysics. We tested with letters, which, after little or no training, provide the many possible targets that are needed for quick testing (Pelli & Robson, 1991). Long term, we are interested in testing crowding in children (Waugh et al., 2018) as an early biomarker for susceptibility to visual problems like dyslexia. Crowding distance is highly conserved across object kind (Kooi et al., 1994; Pelli & Tillman, 2008), which suggests that letters, vernier, and Gabors might have similar crowding distances, but Grainger et al. (2010) reported different crowding distances for letters and symbols.

Crowding exhibits several striking asymmetries. Crowding distance measured radially — along a line passing through the foveal center — is roughly twice that measured tangentially — the orthogonal orientation (Greenwood et al., 2017; Kwon et al., 2014; Pelli, 2008; Petrov & Meleshkevich, 2011; Toet & Levi, 1992). Crowding distance has often been reported to be smaller in the lower than upper visual field (Fortenbaugh et al., 2015; Greenwood et al., 2017; He et al., 1996; Petrov & Meleshkevich, 2011) and on the horizontal than vertical midline (Chung, 2013; Coates et al., 2021; Liu et al., 2009; Petrov & Meleshkevich, 2011; Toet & Levi, 1992; Wallis & Bex, 2012).

Crowding distance is a potentially valuable biomarker for several reasons. Crowding severely limits what we see, how fast we read, and is associated with dyslexia. There are large individual differences in crowding distance and correspondingly large physiological differences in the sizes of relevant areas of visual cortex, which invite analysis by correlation (Kurzawski et al., 2021). Here we measured crowding in 50 observers. Previous in-person crowding surveys (Grainger et al., 2010; Greenwood et al., 2017; Petrov & Meleshkevich, 2011; Toet & Levi, 1992) included at most 27 observers. The only remote crowding survey tested 793 observers but did not report any asymmetries (Li et al., 2020). The above cited works used various kinds of stimuli, including letters of various fonts. The original reports of the crowding phenomenon were mostly letter-based (Anstis, 1974; Bouma, 1970, 1973; Ehlers, 1936; Ehlers, 1953; Korte, 1923; Stuart & Burian, 1962). Historical review of crowding is described elsewhere (Levi, 2008; Pelli et al., 2004; Strasburger, 2020; Strasburger et al., 2011). Here, we too use letters, because they do not require training, and provide a large number of stimulus alternatives, which speeds threshold estimation in laboratory and clinical testing (Pelli et al., 1988).

Whether crowding can be explained by the neural computations in any particular cortical location remains unknown, but several candidate areas have been suggested: V1 (Millin et al., 2014), V2 (Freeman & Simoncelli, 2011; He et al., 1996), V3 (Bi et al., 2009; Tyler & Likova, 2007), hV4 (Burchell et al., 2019; Liu et al., 2009; Motter, 2006; Zhou et al., 2017) and higher-order areas (Aghdaee, 2005; Louie et al., 2007). The magnitude of the BOLD signal in V1 is lower in the presence of crowding (Millin et al., 2014). Crowding distance is different for stimuli tuned to stimulate either the parvo- or magno-cellular pathway (Atilgan et al., 2020). Although both crowding and acuity increase linearly with eccentricity, which might suggest a common physiological origin, the two lines have very different intercepts with the eccentricity axis, i.e. the E_&_ value for acuity is more than 5 times larger than the E_2_ value for crowding E_2_ = 2.72 for acuity and E_2_ = 0.45 for crowding] (Latham & Whitaker, 1996; Petrov & Meleshkevich, 2011; Rosenholtz, 2016; Song et al., 2014; Strasburger, 2020).

This seems inconsistent with a common cause. Here, we use our data from 50 observers to study the relationship between acuity and crowding in the fovea.

In our 50 participants, we measured 13 crowding distances at three eccentricities (0, 5, and 10 deg), on all four cardinal meridians, in two crowding orientations (radial and tangential), using two fonts (Sloan and Pelli). We also measured acuity in the fovea. We used the Sloan and the Pelli fonts. As far as we know, the Pelli font is still the only letter font skinny enough to measure crowding distance in the fovea. Apart from letters, foveal crowding can be measured with vernier targets (Malania et al., 2007). We also assessed crowding’s variation along the four cardinal meridians, in two crowding orientations, and across individuals.

Crowding varies two-fold across meridians, producing several asymmetries.

## METHODS

### Measuring crowding

We measured thresholds of many participants, collecting two datasets with similar methods except for one important difference. Details about Quest and stimulus presentation were similar for both and are described in the sections below. Here we focus on the differences.

*Method 1: Unmonitored fixation*. Threshold was measured without gaze tracking. Viewing distance was measured before each session, and no chin rest or forehead support was provided. The participant identified the target by pressing that letter in the keyboard.

Participants were naive to the task and received no advance training. In each block, crowding distance was measured at two randomly interleaved target locations, which were horizontally or vertically symmetric about the fixation cross. This *unmonitored*-fixation dataset includes one radial crowding distance on each of the four cardinal meridians with the Sloan font for 100 participants.

*Method 2: Awaited fixation*. Each trial began only once the participant had continuously fixated within ±1.5 deg of the crosshair for 250 ms, and we only saved trials in which gaze remained within ±1.5 deg of the crosshair center until stimulus offset. The experimenter was present during data acquisition. Viewing distance was measured at the beginning of each session and maintained by use of a chinrest with forehead support. Participants identified the target by using a mouse to click on one of the letters displayed on the response screen. For each participant, data collection began after a total of 10 correct trials. Crowding distance was measured at four randomly interleaved target locations symmetric about the fixation cross, one on each cardinal meridian. We acquired two thresholds at each location to estimate test-retest reliability. This *awaited*-fixation dataset includes two radial crowding distances on each of the four cardinal meridians with the Sloan font for 50 participants.

Comparing results obtained with the two methods at ±5 deg on the horizontal midline, reveals large differences in the mean and distribution of the Bouma factor (**Fig. 1**). [In this paper, “log” is the logarithm base 10.] Using *unmonitored* fixation, the geometric mean *b* was 0.12, with a 0.31 SD of log *b*. Using *awaited* fixation, the geometric mean *b* was higher, 0.20, with a lower SD of log *b*, 0.18. The awaited fixation histogram (red) is compact. The unmonitored fixation histogram (green) is much broader, extending to much lower values of *b*. Our interpretation of the broader histogram and lower geometric-mean*b* in *unmonitored* fixation is that observers occasionally “peek”, that is fixate near an anticipated location of the target instead of the fixation cross as instructed. Indeed, at the end of Results, we present a quantitative peeking model showing that peeking reduces the geometric mean*b* and broadens its distribution, consistent with the observed results.

**Figure 1.**
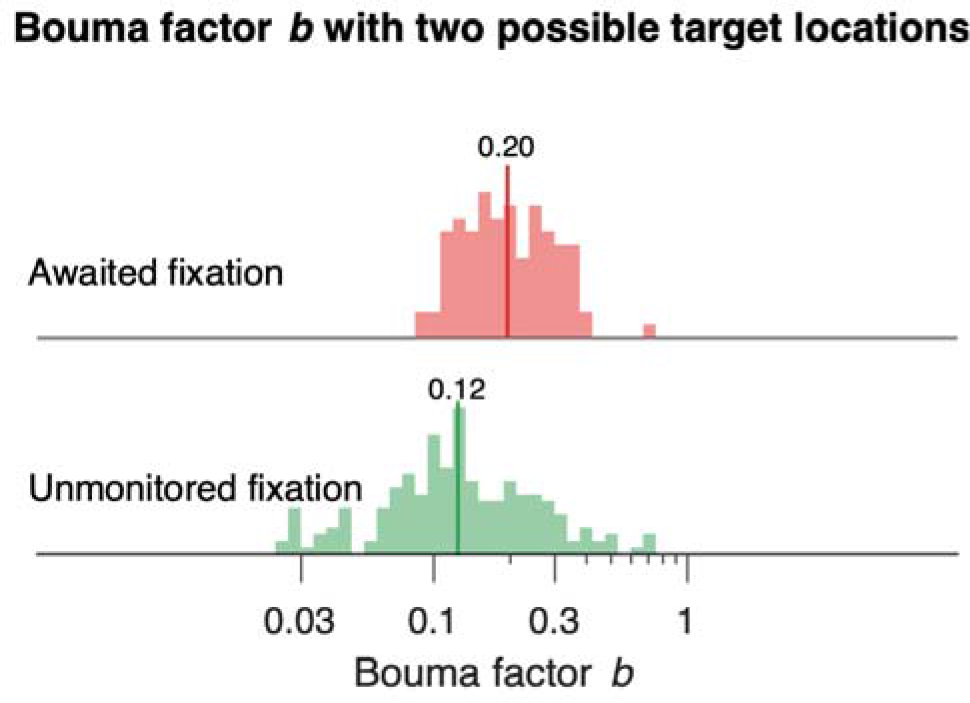
Histograms of the Bouma factor estimated by two methods . Each histogram shows the Bouma factor b at ±5 deg eccentricity along the horizontal midline. For awaited fixation we only used data from first session of the experiment. The geometric mean b is indicated by a dark vertical line capped by a number. *Unmonitored* fixation gave a 0.12 geometric mean *b* with 0.31 SD of log *b*. *Awaited* fixation gave a higher 0.20 geometric mean *b* with a lower 0.18 SD of log *b* (see Table 3). Each histogram includes *b* estimates made at both +5 and -5 deg eccentricity on the horizontal midline. The Results section below reports a 0.78:1 right:left advantage. Note: Mixing data from the two locations (-5 deg left and +5 deg right) makes the combined histogram slightly broader than that for either location. The 0.78:1 *b* ratio corresponds to a -0.107 log *b* difference. If we suppose that mixing log *b* estimates from the two locations is equivalent to taking all the data from the right location and adding +0.107 to a random half of the log *b* estimates, then mixing the two locations increases the variance by +0.0025, which is only 8% of the measured variance of log *b* for awaited fixation, and only 3% for unmonitored fixation.

This paper focusses on the *awaited*-fixation data, which can be downloaded from OSF (https://osf.io/83p6u/).

### Crowding dataset

Data were acquired with the *CriticalSpacing* software (Pelli et al., 2016) using Quest (Watson & Pelli, 1983), allowing for reliable and relatively fast measurement of crowding distances. Our crowding database consists of measurements of crowding distance with the Sloan font (with radial and tangential flankers) and with the Pelli font (radial flankers) in 50 observers. With the Sloan font, we measured crowding at 8 different peripheral locations in the visual field: 2 eccentricities (5 and 10 deg) along the four cardinal meridians (upper, lower, left, and right). Sloan tangential crowding was measured only at ±5 deg eccentricity on the horizontal midline. With the Pelli font, we measured crowding at the fovea and at ±5 deg on the horizontal midline. The Sloan font acuity size is too big to allow measuring foveal crowding distance in adults. The Pelli font was specially designed for measuring foveal crowding distance (Pelli et al., 2016). We also measured acuity in the fovea. A spatial map of the testing is shown below, in **Figure 4** in Results. We also tested 10 observers at 20 and 30 deg eccentricity with radial flankers (only 1 session) and plotted the results in **Figure 9**. To estimate test-retest reliability of our measurements we used two sessions to measure each threshold twice. Sessions were scheduled at least a day apart over a maximum of 5 days apart. We report our results as the Bouma factor *b* (slope of crowding distance vs. eccentricity) estimated from **Eq. 10**, to minimize error in fitting log s^. Here, crowding distance s^ is the required center-to-center spacing (in deg) for 70% correct report of the middle letter in a triplet.

### Participants

**Table 1** describes our main dataset, and **Figure 4** (in Results) plots its spatial coverage of the visual field. The study tested 50 observers (mean age = 23), mostly New York University undergraduate students. Each observer had normal or corrected-to-normal vision. All experiments were conducted in accordance with the Declaration of Helsinki and were approved by New York University’s ethics committee on activities involving human observers. In all analyses, except the test-retest section, we average the first- and second- session thresholds.

**Table 1.** Data summary. In the periphery, we measured crowding distance radially and tangentially with the Sloan font. With the Pelli font, we measured crowding both in the fovea and periphery. We also measured foveal acuity with the Sloan font. Each threshold was measured once in two sessions separated by at least 24 hours. For peripheral thresholds, we used gaze tracking to guarantee fixation within ±1.5 deg of the crosshair center. Foveal crowding required long viewing distance which made gaze tracking impractical, so participants were merely instructed to fixate the center of the crosshair. We suppose good fixation of the central crosshair because the participants expected a foveal target.

| *N* | Measure | Font | Crowding orientation | Radial eccentricity (deg) | Cardinal meridians | Thresholds per observer (each measured twice) | Gaze tracking |
| --- | --- | --- | --- | --- | --- | --- | --- |
| 50 | Crowding | Sloan | radial | 5, 10 | all | 8 | Yes |
|  | Crowding | Sloan | tangential | 5 | right, left | 2 | Yes |
|  | Crowding | Pelli | radial | 5 | right, left | 2 | Yes |
|  | Crowding | Pelli | horizontal | 0 | — | 2 | No |
|  | Acuity | Sloan | — | 0 | — | 2 | No |

The peeking-model section in Results refers to these “awaited-fixation” peripheral measurements on 50 participants and compares them to separate “unmonitored fixation” peripheral measurements on 100 participants.

### Apparatus

Each testing session was completed on an Apple iMac 27” with an external monitor. The observer viewed the LG 27” 5K monitor 27MD5KL-B, with a screen resolution of 5120 x 2880 and a white background with luminance 275 cd/m2. The white background never changed throughout the experiment; the black crosshair and letters were drawn on it. The observer viewed the screen binocularly at one of several different viewing distances. The software required a special keypress by the experimenter at the beginning of every block with a new observer or a new viewing distance, to affirm that the new viewing distance (eye to screen) was correct as measured with a tape measure, and that the screen center was orthogonal to the observer’s line of sight. To measure crowding and acuity in the fovea, the viewing distance was 200 cm. For ±5 and ±10 deg eccentricity the distance was 40 cm, and for ±20 and ±30 deg it was 20 cm. The long viewing distance gives good rendering of small stimuli; the short viewing distance results in a wide angular subtense of the display, to allow presentation of peripheral targets on either side of a central fixation.Stimuli were rendered using *CriticalSpacing.m* software (Pelli et al., 2016) implemented in MATLAB 2021 using the Psychtoolbox (Brainard, 1997; Pelli, 1997). Every Sloan letter was at least 8 pixels wide and every Pelli digit was at least 4 pixels wide.

### Stimuli and procedure

To measure acuity, we show one letter. To measure crowding we show a trigram of three letters or digits. For each trial, the three letters or digits are drawn randomly, without replacement, from the 9 letters (DHKNORSVZ) or digits (123456789) available. Letters and digits are rendered as black in the Sloan or Pelli font, presented on a uniform white background (Pelli et al., 2016; Sloan et al., 1952). We omit the C in the Sloan font because it’s too easily confused with the O (Elliott et al., 1990). For crowding, each trigram was arranged either radially or tangentially. Each testing session included several blocks and was about an hour long. Most blocks measured 4 thresholds, interleaved, usually four crowding thresholds on the 4 cardinal meridians at the same radial eccentricity. For the Pelli font and tangential crowding we measured two thresholds at symmetric locations about fixation along the horizontal midline. To minimize the temptation to look away from fixation toward an expected target location, we randomly interleaved conditions measuring threshold at the same radial eccentricity at 2 or 4 symmetric locations around fixation. A sample stimulus sequence appears in **Figure 2A**. A central crosshair (the fixation mark) is displayed until the observer presses a key to initiate the trial. Then, after 250 ms of correct fixation, the letter trigram appears on the screen for 150 ms and the computer waits for the observer to identify the middle stimulus letter by using a mouse to click on a letter in a row of all the possible letters on the response screen. Observers are instructed to return their eyes to fixation before clicking their response. A correct response is acknowledged with a brief beep. Then the computer waits indefinitely for the observer to fixate within 1.5 deg of the crosshair for 250 ms, and then immediately presents the stimulus for the next trial. If the observer fails to fixate for 250 ms within a 10 second window, the software asks for recalibration of the gaze tracker.

**Figure 2.**
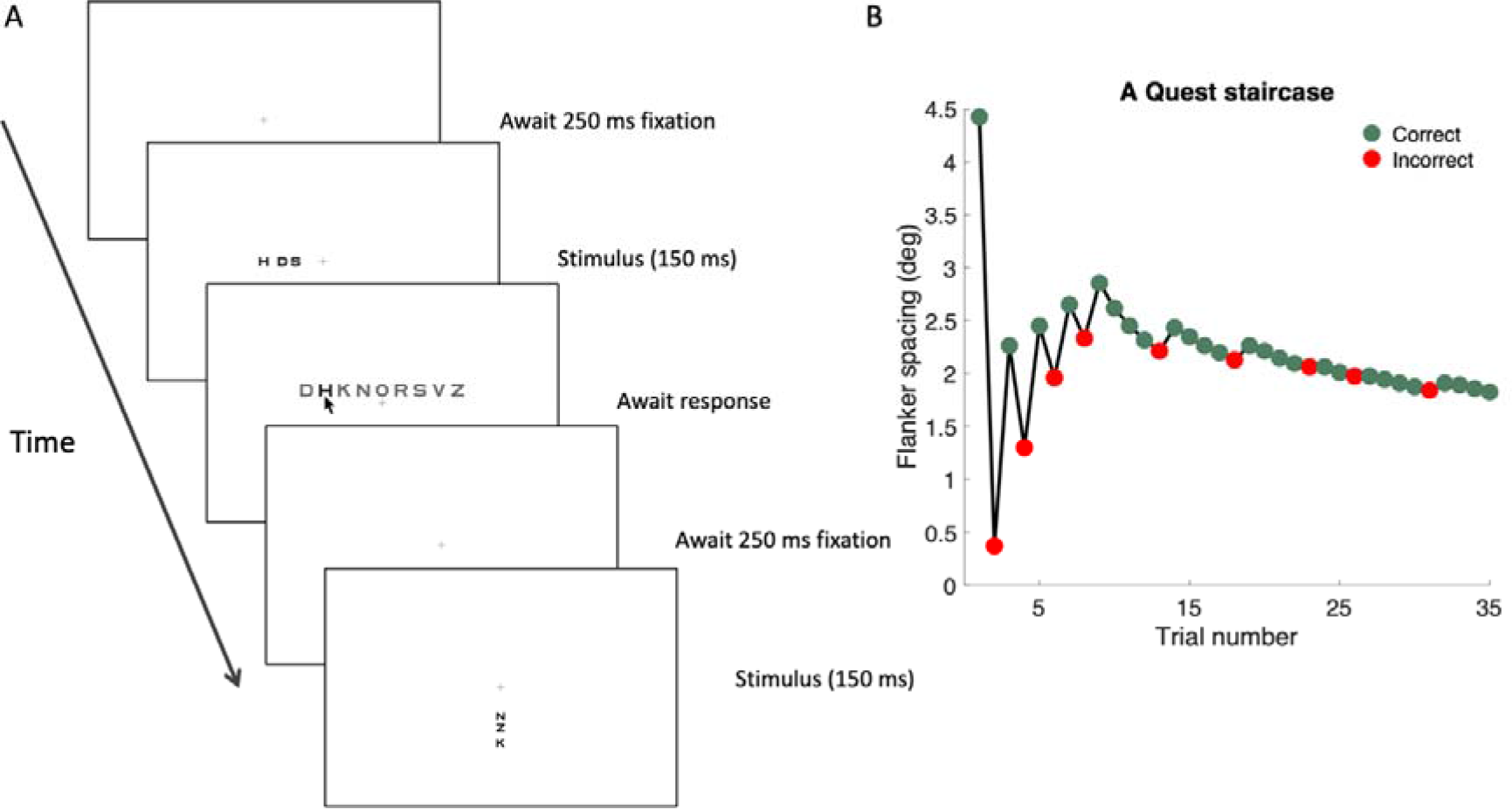
Stimulus and procedure. A) The display sequence for a peripheral trial and part of the next. While gazing at the crosshair, which is always present, the observer presses a space bar key, which begins the first trial. The target is presented only if the observer was continuously fixating for 250 ms. Stimulus presentation is accompanied by a low-pitched purr. Then the observer identifies the target by using a mouse to click on one letter out of all possible letters that appear above the fixation on the response screen. If the response was correct, the observer hears a brief beep acknowledging correctness and silence otherwise. Then the computer again waits for 250 ms of fixation within 1.5 deg of the crosshair center. The four conditions (one for each meridian) are randomly interleaved, so the observer does not know which location comes next. **B)** The staircase sequence of spacings tested on 35 successive trials of one condition (+5 deg eccentricity), under control of QUEST. On each trial, the letter size was a fraction 1/1.4 of the spacing. QUEST picks the most informative spacing to test on each trial to minimize variance of its final threshold estimate. Finally, after 35 trials, QUEST estimates the crowding distance, i.e., spacing to achieve 70% correct. Notice that testing quickly homes in on threshold.

### Measuring crowding distance

Crowding distance was estimated using the Pelli et al. (2016) procedure. Letter spacing is controlled by QUEST. Spacing scales with letter size, maintaining a fixed ratio of 1.4:1. We set the Weibull function guessing rate parameter y to the reciprocal of the number of characters in the test alphabet for that font, usually 9. We set the “finger error” probability o to 1%, to help QUEST cope with an occasional reporting mistake. We set the Weibull function steepness parameter f to 2.3, based on fits to two observers’ psychometric data for radial crowding. At the end of each block, QUEST estimates threshold (crowding distance in deg, **Fig. 2B**). To measure acuity, we followed a similar procedure, except that the target was presented without flankers. Threshold was defined as the letter spacing (crowding distance) in deg or letter size (acuity) in deg that achieved 70% correct identification, using Quest to control the stimulus parameters trial by trial, and make the final estimate.

Each threshold measurement was based on 35 trials (one condition). A block consists of all the trials in however many conditions are randomly interleaved, e.g., 4 × 35 = 140 trials to measure threshold at 4 meridians. Each condition measures threshold for one meridian. The interleaving keeps the observer uncertain as to which location is tested on each trial. We do this to minimize the urge to “peek” away from fixation. On a crowding trial, until the target and flankers appear there is nothing about the display that distinguishes which of the interleaved conditions this trial belongs to.

### Gaze tracking

We used gaze-contingent display to guarantee fixation while measuring all peripheral thresholds. We used an EyeLink 1000 eye tracker (SR Research, Ottawa, Ontario, Canada) with a 1000 Hz sampling rate. To allow short viewing distance (40 cm) we used the EyeLink Tower mount with a 25 mm lens mounted on the EyeLink camera. Each trial presented the stimulus once gaze had been within 1.5 deg of the crosshair center (the fixation mark) for 250 ms. If, during the stimulus presentation, gaze deviated more than 1.5 deg from the crosshair center, then the trial was not saved, the fixation cross turned red (to alert the participant), and the trial was repeated with a fresh letter trigram. Thus, each threshold estimate was based on 35 trials with fixation within 1.5 deg of the crosshair center. The foveal thresholds demanded a long 200 cm viewing distance that was incompatible with our gaze tracking set up, so they were measured without gaze tracking, but fixation is generally good when the participant knows that the target is foveal.

### Model fitting

It is generally found that SD of repeated measurement of threshold spacing*s* is roughly proportional to mean spacing, but the SD of log spacing *S* is independent of mean spacing. Therefore, our fitting minimizes the RMS error in log spacing *S* = log_10_ *s*. The fitting is nonlinear (using the MATLAB fmincon function) because we minimize error in *S*, whereas each model is linear in *s*, not *S*. We estimate the participant, meridional, crowding orientation, and font factors by solving several models (see model equations in Table 4).

Our fitting minimizes the RMS error in predicting the log crowding distances, which is equivalent to minimizing the summed square error, *SSE*:

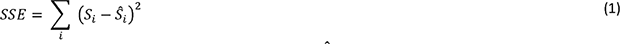

where 5_$_ is the *i*-th log crowding distance, and 5^S^_$_ is the *i*-th predicted log crowding distance. The variance explained by each model is:

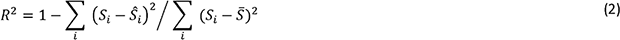

where *S* = mean_i_ *S*_i_ is the mean log crowding distance.

### Model comparison

An F test was used for pairwise model comparison. The model with fewer parameters is called “simple”, and the model with more parameters is called “full”. After calculating the sum of square errors *SSE*_simple_ and *SSE*_full_ for each model (**Eq. 1**), we calculate the F-statistic:

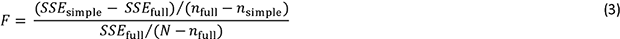

Where *n*_full_ is the number of parameters in the full model, *n*_simple_ is the number of parameters in the simple model, and *N* is the number of observations. We estimate the p-value using the F-distribution. A p-value less than 0.05 indicates that the model with more parameters provides a significantly better explanation of the data.

### Cross validation

First, we divide the thresholds into 6 random subsets of equal size. In each cross-validation step, one subset of data is retained as the validation set for testing the model, while the remaining subsets are used as training data. Each subset is chosen only once for testing. We repeat leave-one-out testing 6 times to obtain the full dataset. Variance explained*R*^2^ is calculated by**Eq. 2.**

### Standardized Bouma factor can be compared across studies

To facilitate comparison across studies and the cooperative evaluation of the Bouma factor as a biomarker of visual health, we define the *standardized* Bouma factor *b*’ as the slope of crowding distance vs radial eccentricity multiplied by a correction factor that accounts for methodological differences from Bouma’s number of choices (25), threshold criterion (75% correct), and linear spacing (vs. log).

**Table 2** computes the correction factors needed to compare the Bouma factor *b* across studies that used various numbers of response choices (e.g., 9 Sloan letters or 2 orientations of a tumbling T), various threshold criteria (e.g., 70% or 75% correct), and linear or log flanker spacing. Including the present one, we know of five studies that compare crowding distance across meridians. We take Bouma (1970) as the standard for this standardized way of reporting the strength of crowding.

**Table 2.**
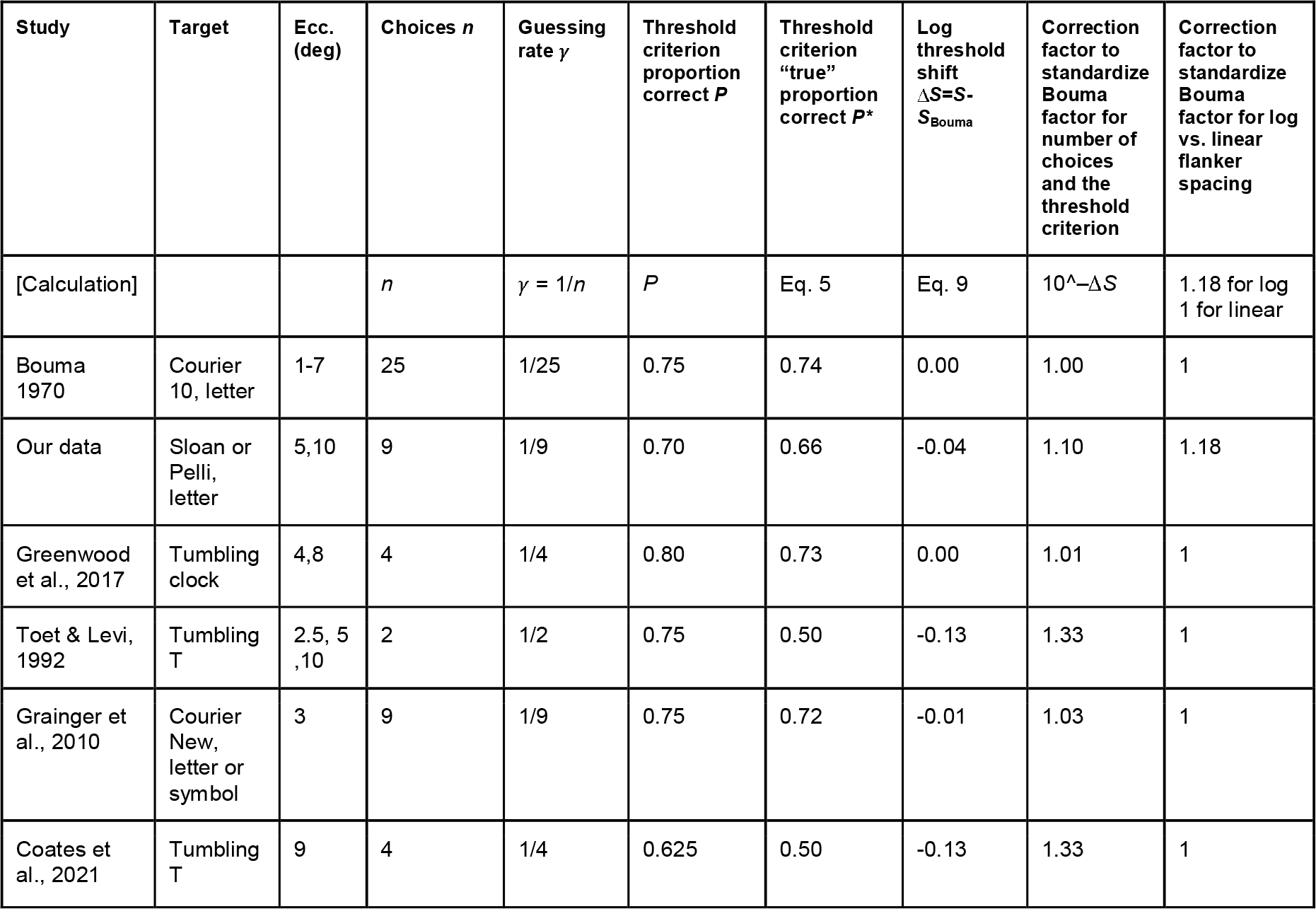
Correction factor to standardize each study’s Bouma factor to compare studies. The resulting standardized Bouma factors are reported in Table 6. Each study’s difference from Bouma’s 25 choice alternatives and 75% correct threshold criterion offset the threshold log spacing by *ΔS* relative to Bouma’s. The correction factor is 10^– *ΔS*. We provide a brief Excel spreadsheet to calculate the correction factor and a MATLAB script that produces plots like Fig. 3. The correction factor to account for linear vs. log flanker spacing is estimated in the Supplementary material. Note that crowding measured with tangential flankers does not require correction for log vs. linear spacing symmetry. Tangential spacing was always linear.

The log-threshold shift *ΔS* is illustrated in **Figure 3B**. Using the Coates et al. (2021) reanalysis of Bouma’s (1970) data, we estimated the Bouma factor for a 75% threshold criterion applied to Bouma’s (1970) results (see the Bouma factor paragraph in Discussion).

**Figure 3.**
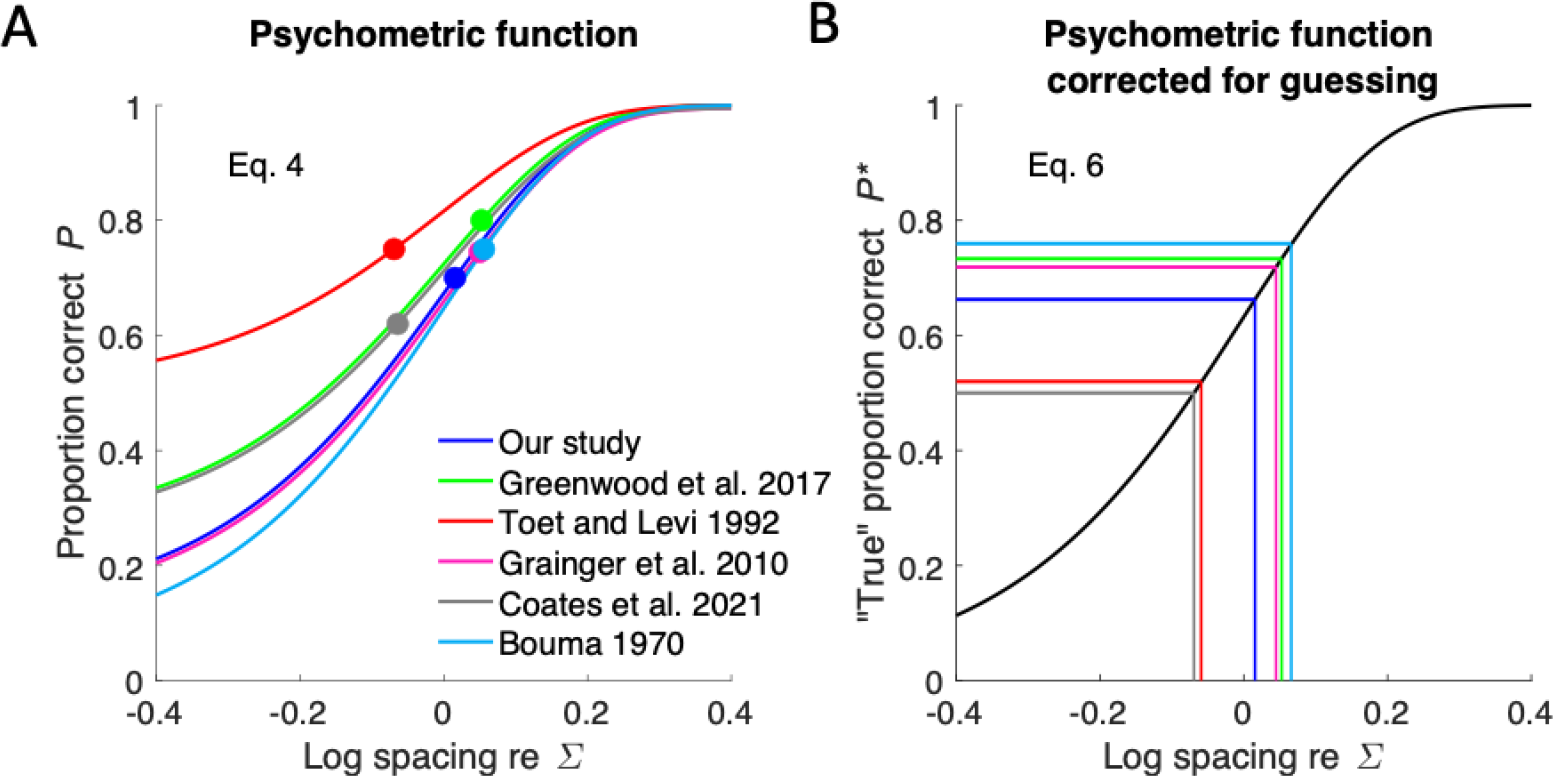
Effect of guessing rate and criterion on threshold log crowding distance. S**. A) Proportion correct (Eq. 4)** of the six studies, with threshold parameter J; set to zero, showing the effects of the number of response choices *n* (sets lower asymptote γ= 1/*n*) and the threshold criterion (height of each colored dot). **B) “True” proportion correct (Eq. 6)** for the same studies. Each study’s threshold criterion *P\** is represented by a horizontal line. For each study, a vertical line reads off the log threshold spacing *S* at its threshold criterion *P\**. Results from Bouma (1970) are used with the Andriessen and Bouma (1976) 75% threshold criterion. Table 2 computes the difference between each study’s threshold and Bouma’s. To avoid occlusion, the Toet and Levi and Bouma lines in this panel were offset by +0.02 horizontally and vertically.

**Figure 4.**
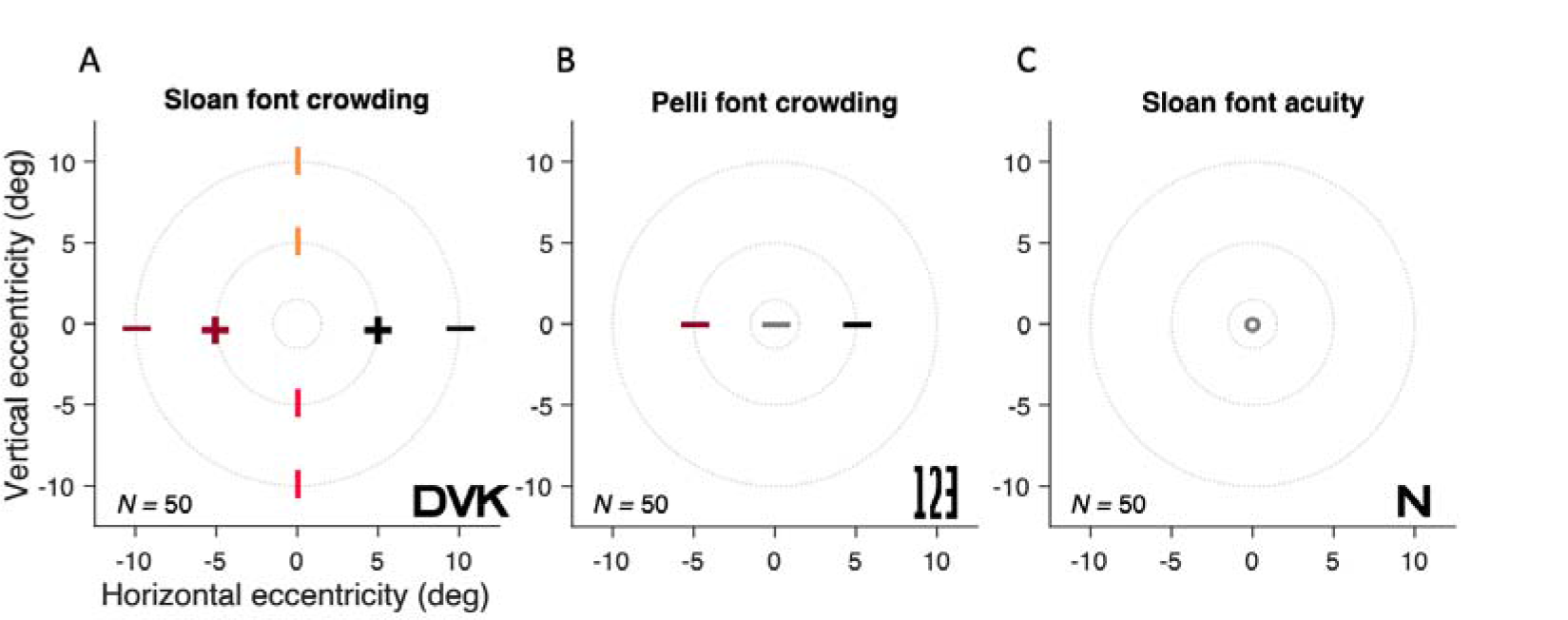
**Map of testing**. Each panel title indicates the font and threshold task. The number of observers tested is indicated in the lower left. The +, –, and o symbols indicate testing of crowding with radial (—) or radial and tangential (+) flankers, and testing of acuity (o). Typical stimuli appear in the lower right of each panel. Beyond the main dataset described here, Fig. 9 shows additional results from 10 observers at 20 and 30 deg eccentricity.

**Figure 5.**
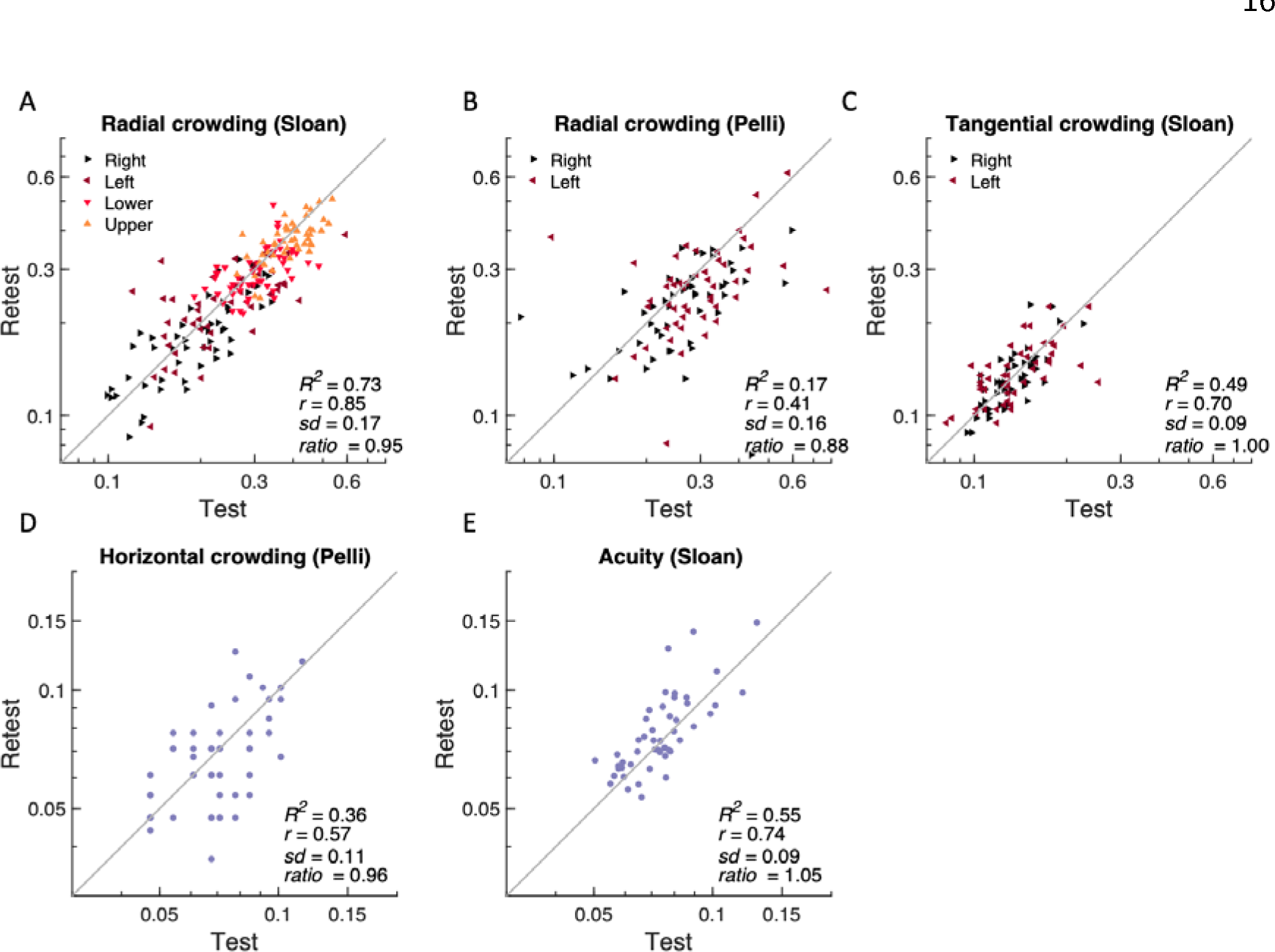
Test-retest reliability of threshold estimates. . Estimates of Pearson’s *r* correlation coefficient, standard deviation, retest:test ratio, and *R*^2^ are based on the log of the font-task threshold, in degrees of visual angle, named at the top of each panel. For peripheral crowding (**A-C**) each measurement is represented by a triangle pointing towards the tested meridian. The gray line equality.

*Proportion correct.* To account for the different number of choices and the threshold criterion, we assume that, with accurate fixation, the proportion correct *P* is a Weibull function of the log spacing *S*,

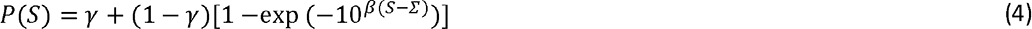

with a threshold parameter *Σ,* where *γ* is the guessing rate (the reciprocal of the number of target choices, which is 1/25=0.04 in Bouma’s 1970 results), and *β* is the steepness parameter, which we set to 2.3, based on fitting psychometric functions to hundreds of trials at several spacings by two experienced observers. **Figure 3A** shows this psychometric function for six studies, taking the guessing rate γ to be the reciprocal of the number of choices *n*, and using our own estimate of the steepness parameter *β*=2.3 .

*“True” proportion correct*. To accommodate various numbers of choices *n*, and thus guessing rates *γ*=1/*n*, we “correct for guessing,”

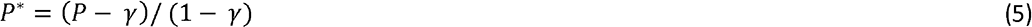

This is a popular transformation of psychometric data, usually justified by assuming that the guessing rate can be modelled as an independent process. Because it discounts false alarms, the corrected hit rate is called the “true hit rate”. That makes sense for a yes-no task, but not for an identification task. Here we proceed regardless, and compute the “true” proportion correct, because, with this Weibull function (**Eq. 4**), correction for guessing (**Eq. 5**) removes all dependence on *γ*. Applying correction for guessing to any given threshold criterion *P* gives us the corresponding “true” proportion correct criterion *P\** to apply after correction for guessing. Similarly applying correction for guessing to the psychometric function (**Eq. 4**) gives us the “true” proportion correct,

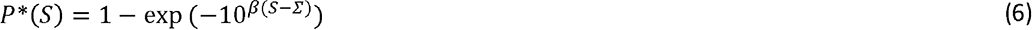

The inverse of **Eq. 6** is

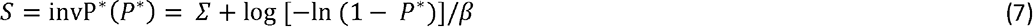

**Figure 3B** plots “true” proportion correct (**Eq. 6** with *β*=2.3 and *Σ*=0), the same function for all studies, and, for each study, a vertical line reads off the log threshold spacing *S* at its threshold criterion *P\**.

*Relative to the Bouma standard.* Thus, a study’s number of choices and threshold criterion increase its log threshold by Δ*S* relative to the Bouma standard:

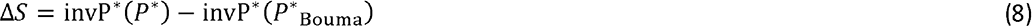

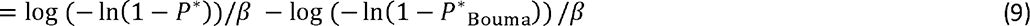

where *β*=2.3, and *P*\* and *P*\*_Bouma_ are the “true” proportion correct threshold criteria computed by**Eq. 5** from the study’s criterion *P* and Bouma’s *P*_Bouma_=0.75 (Andriessen & Bouma, 1976)

### Log-symmetric spacing of flankers

Since (Bouma, 1970), most crowding studies measure crowding distance as the center-to-center spacing between the target and each of two flankers on opposite sides of the target that yields a criterion level of performance. When crowding is measured in the radial orientation, the Bouma law tells us that crowding distance increases linearly with eccentricity. Several studies have documented that, when flankers are arranged symmetrically about the target on a radial line from fixation, the outer flanker has much more effect (Banks et al., 1977; Bex & Dakin, 2005; Estes et al., 1976; Krumhansl, 1977). This is to be expected since crowding distance grows with eccentricity and the outer flanker is more eccentric. In fact, crowding distance on the cortical surface (in mm)—the product of crowding distance in deg and cortical magnification in mm/deg—is conserved across eccentricity (for eccentricities above 5 deg) because psychophysical crowding distance scales with eccentricity (Bouma, 1970; Kooi et al., 1994; Levi & Carney, 2009; Pelli et al., 2004; Toet & Levi, 1992). Given the logarithmic cortical mapping of the visual field (Fischer, 1973), when measuring radial crowding we space the trigram so that the log eccentricity of the target is midway between the log eccentricities of the flankers and report the inner spacing. This raises the question of how to compare crowding distances between experiments that spaced the flankers linearly vs logarithmically. Given the Bouma law (**Eq. 8** below), supposing that crowding distance depends primarily on the flanker-to-flanker distance and only negligibly on the target position between them, we show in the Supplement (“Effect of symmetric placement of flankers with regard to either linear or log eccentricity”) that the crowding distance is expected to be 1.18 times larger when measured with linearly-spaced flankers than with log-spaced flankers. To ease comparison across studies, the correction factors in Table 2 include this effect of log vs linear spacing on the estimated Bouma factor.

### Might attention help explain differences in the reported Bouma factor?

Attention reduces many perceptual thresholds (Carrasco, 2011). Many researchers have assessed the effects of attention on crowding, but they have yet to reach a consensus. Several found an attentional benefit in crowding tasks (Bacigalupo & Luck, 2015; Kewan- Khalayly et al., 2022) including reduction of crowding distance (Yeshurun & Rashal, 2010), but others did not find such effects (Scolari et al., 2007; Strasburger, 2005; Strasburger & Malania, 2013). All our peripheral crowding thresholds are measured with either two-fold or four-fold uncertainty about target location, and we suppose that attention was distributed among the possible target locations. It is possible that attentional bias contributes to some of the Bouma factor asymmetries.

### Data from other studies

Data were extracted from Figure 6 of Toet and Levi (1992) using WebPlotDigitizer (Rohatgi, 2020). Data were extracted from Figure 7 of Grainger et al. (2010). Data from of Greenwood et al. (2017) Supplementary Fig. 1 and Coates et al. (2021) Figure 10 were received as personal communications from the authors. Data from Bouma (1970) were used by means of the recent reanalysis by Coates et al. (2021).

**Figure 6.**
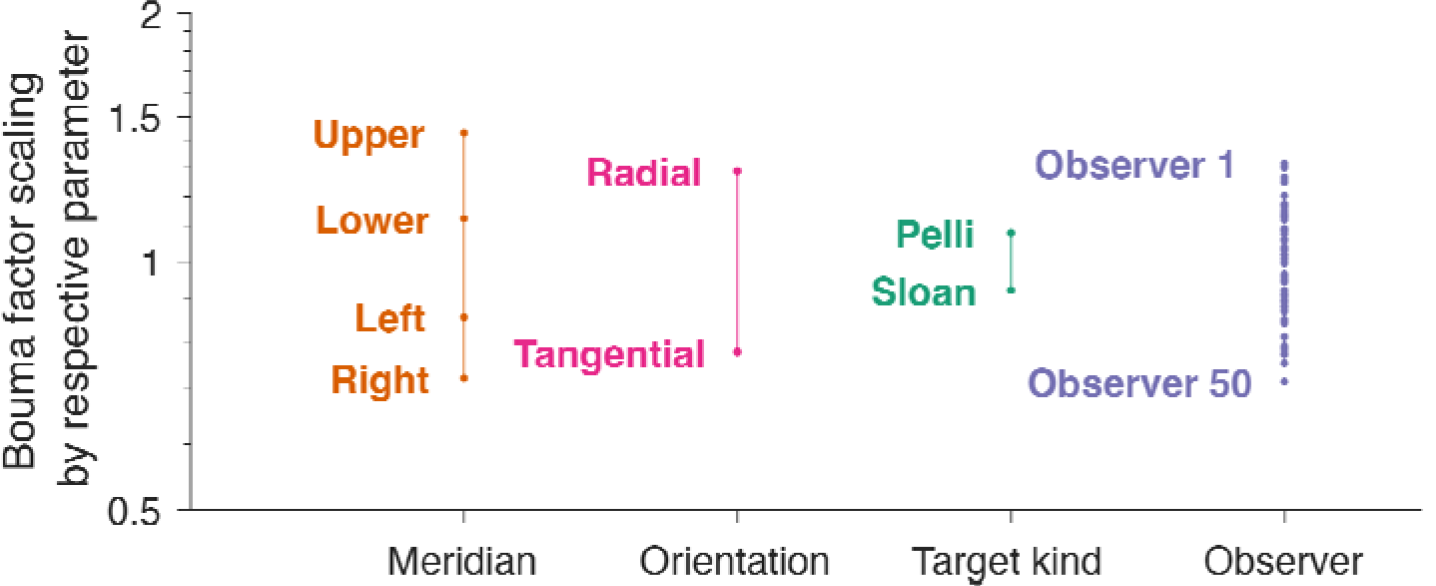
How several parameters scale the Bouma factor. To reveal the effect of each parameter (horizontal axis) each set of model parameters was normalized by the geometric mean of that set. The vertical axis plots the model’s estimates of that parameter. Because the model is multiplicative, the final Bouma factor is proportional to the product of all the parameters. We find a similar variation of Bouma factor with meridian, crowding orientation, and observer. Target kind has the least effect, but that is partly because nearly all of our data are with one font.

**Figure 7.**
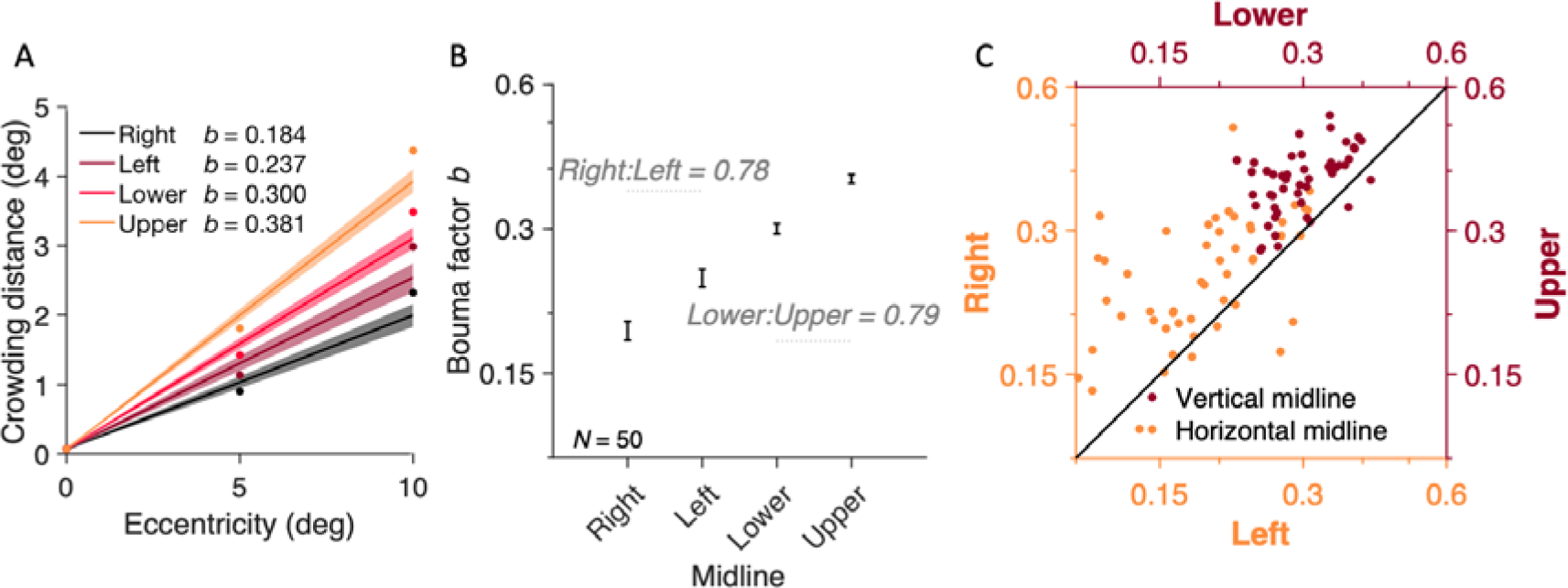
Bouma factor vs. meridian. A) Bouma law estimates for radial crowding with the Sloan font estimated using **Eq. 10**. Each point represents the mean across participants and error bars represent 95% confidence intervals. **B)** Bouma factor vs. meridian. **C)** Individual-participant data plotted for the vertical (dark red) and horizontal midline (orange).

**Figure 8.**
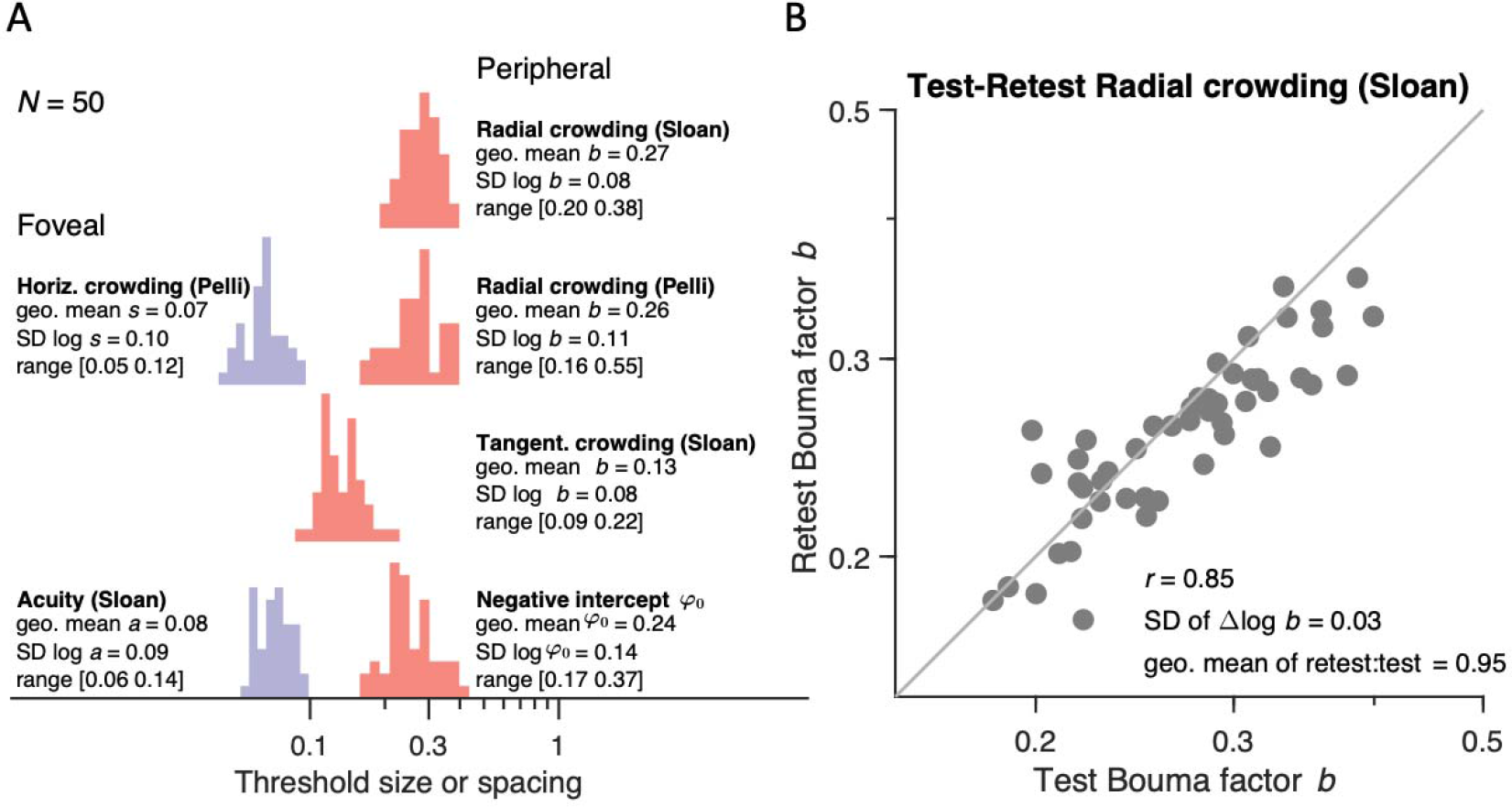
Histograms of crowding and acuity. Histograms of **A)** radial Sloan, radial Pelli, and tangential Sloan Bouma factor (dimensionless), and negative intercept *φ*_0_ (in deg), horizontal crowding distance *s* (in deg), and acuity *a* (in deg). To estimate individual differences, we used all acquired data (e.g., 16 thresholds for radial crowding with Sloan font or 4 thresholds for radial crowding with Pelli font). **B)** Retest vs test of Bouma factor *b* for radial Sloan.

**Figure 9.**
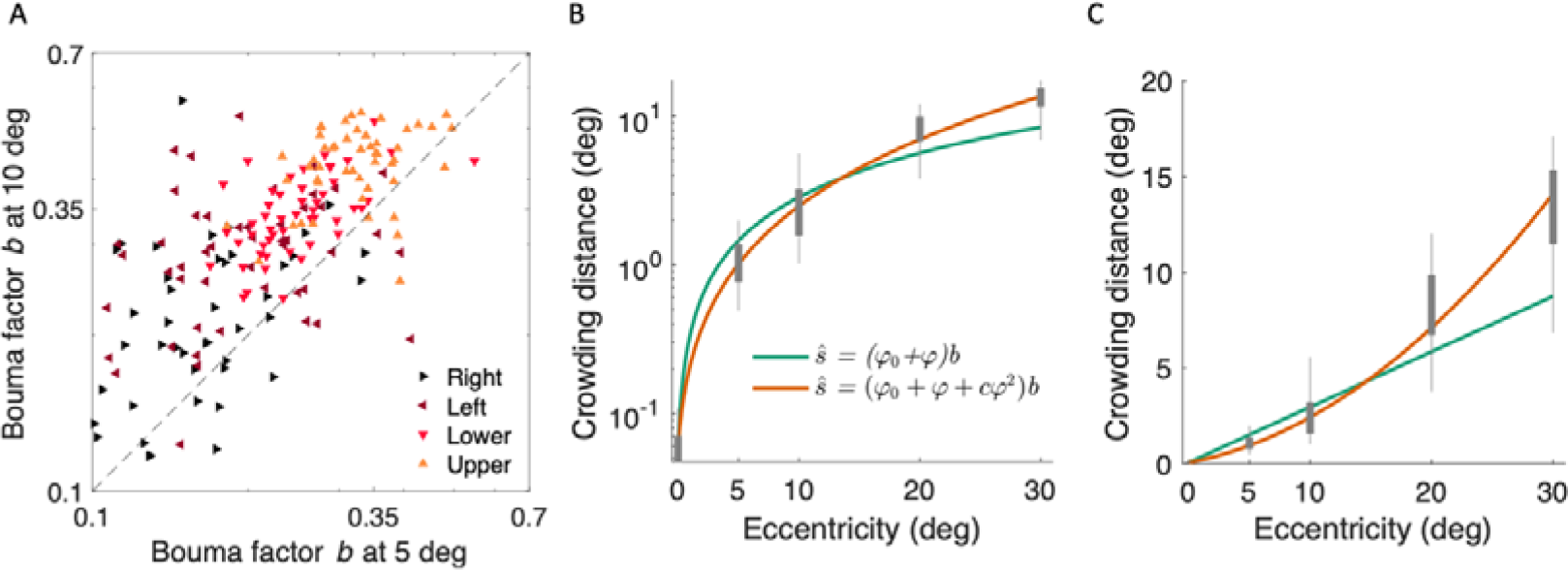
Supralinearity: Crowding distance slope grows with eccentricity. A) Radial Bouma factor at 5 vs. 10 deg eccentricity for Sloan font. Each color shows data on a different meridian. Data are plotted for all 50 participants included in the main study. **B)** Log crowding distance for 10 participants is plotted against eccentricity out to 30 deg. The linear Bouma law is green, and the quadratic Bouma law is orange. We use log coordinates because the fit minimizes the error in log coordinates. Enhancing the Bouma law from linear to quadratic increases the explained variance from 90% to 95%. **C)** Same fits replotted in linear coordinates. The nonlinear growth of crowding distance with eccentricity is not an artifact of perspective transformation: The computation of target angular size and eccentricity was done correctly using the arc tangent function. Eq. 10 fit with RMSE 0.20 and *b =* 0.30, and *φ*_0_ = 0.20. **Eq. 15** fit with RMSE 0.16 and *b =* 0.15, *φ*_0_ = 0.43, and *c* = 0.06.

**Figure 10.**
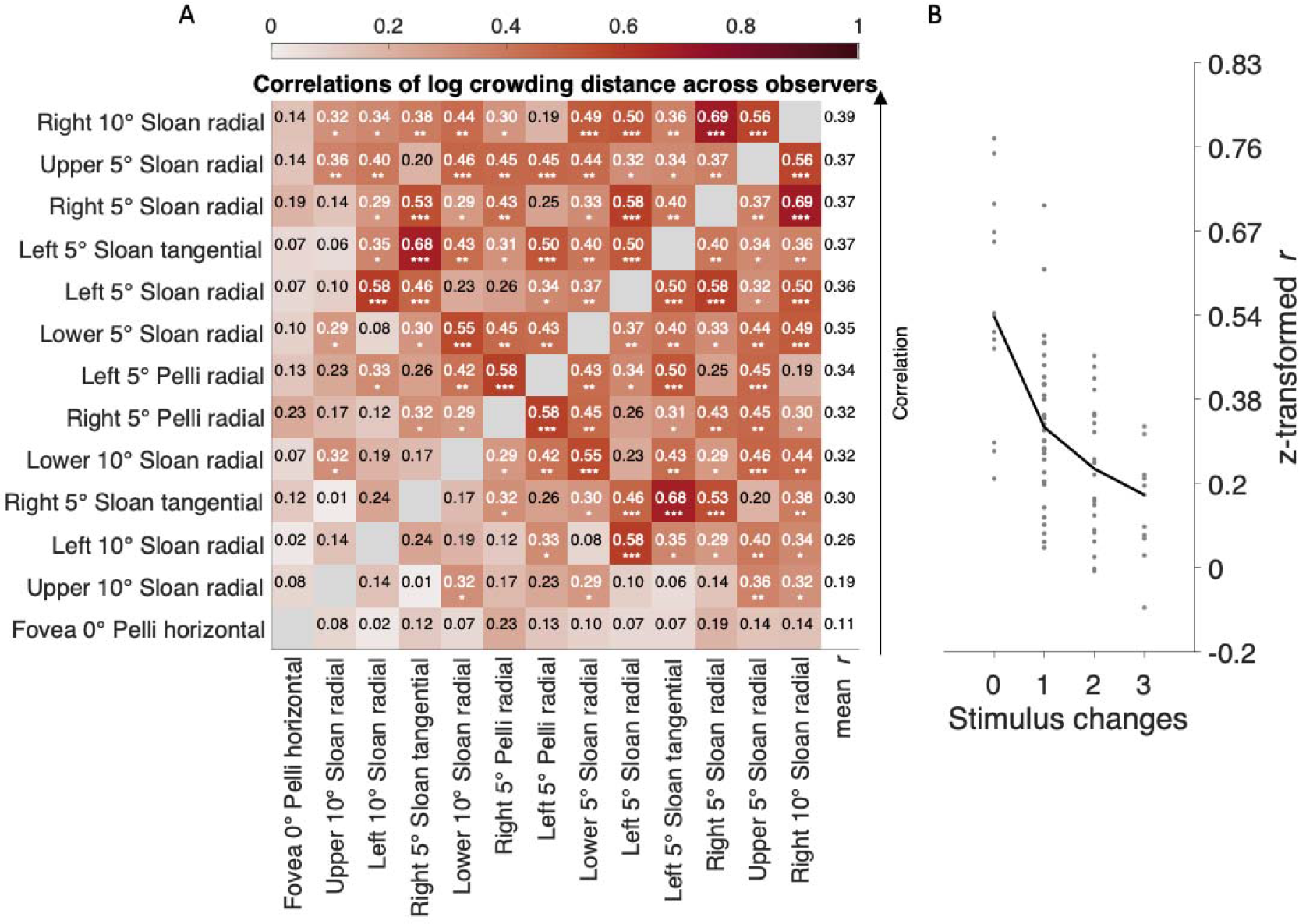
Correlations of log crowding distance. A) Pair-wise correlations between crowding distances for various conditions (Table 1). Rows are sorted so that the average correlation decreases from top to bottom. **B)** Average correlation as a function of the number of stimulus property differences in: radial eccentricity, meridian, crowding orientation, and target kind. For example, comparing two eccentricities is a 1-parameter difference.

### Statistical analysis

Statistics of the log Bouma factor *B* = log *b* were assessed by an ANOVA with *B* as the dependent variable. Two sample comparisons are made with the Wilcoxon rank sum test. We report Pearson’s *r* correlation coefficient for test-retest reliability and correlations of crowding distance.

## RESULTS

### Crowding and acuity

As shown in the map of testing (**Fig. 4**), radial crowding thresholds were measured in 50 adults at 9 visual field locations. Using the Sloan font, radial crowding thresholds were measured at the four cardinal meridians at 5 and 10 deg eccentricity, and tangential crowding thresholds were measured on the left and right meridians at 5 deg eccentricity. Using the Pelli font, the horizontal crowding threshold was measured, in the fovea and on the right and left meridians at 5 deg eccentricity. Foveal acuity was also measured with the Sloan font. Each threshold was measured twice.

### Test-retest reliability of visual threshold

Measurement reliability was assessed by measuring each threshold twice, at least one and no more than five days apart. Crowding thresholds are converted to Bouma factors*b* (see **Eq. 10** below). Foveal crowding and acuity are presented as crowding distance (deg) and acuity as letter size (deg). **Figure 5** plots a scatter diagram of estimates from first vs second session for each combination of font and task. For the Pelli font (**Fig. 5B**), we found a clear improvement of measured crowding distance in the second session (ratio of geometric mean retest:retest = 0.88). This training benefit was much smaller for the Sloan font (0.95), presumably because Sloan is more similar (than Pelli) to familiar fonts. In general, each threshold is derived from a Quest staircase with 35 trials, which takes about 3.5 minutes, and has very good reproducibility. The analyses performed in the following sections are based on the geometric average threshold across both sessions.

### Analysis of variance

**Table 3** presents an ANOVA analysis of the radial Sloan Bouma factors (also plotted in Figure 5A). The 0.18 SD for radial Sloan with 2 meridians (with awaited fixation) in**Figure 1** corresponds to the 0.17 total SD with 4 meridians (also with awaited fixation) in**Table 3**.

**Table 3.** – Analysis of variance. We computed the contribution of each parameter to overall variance. Meridian contributes the most (SD = 0.12) and test-retest contributes the least (SD = 0.01). Degrees of freedom d.f. is number of parameters minus 1. Error is the remaining variance not accounted for by a linear combination of meridian, observer, and test-retest. There were no significant pairwise interactions among meridian, test-retest, and observer (all *p*>0.5).

| Factor | d.f. | Variance | SD | p-value |
| --- | --- | --- | --- | --- |
| meridian | 3 | 0.0143 | 0.1195 | <0.01 |
| observer | 1 | 0.0059 | 0.0770 | <0.01 |
| test-retest | 59 | 0.0001 | 0.0109 | 0.016 |
| error | 346 | 0.0071 | 0.0843 |  |
| total | 399 | 0.0274 | 0.1656 |  |

The 0.08 SD for Sloan radial crowding in **Figure 8** corresponds to the 0.08 SD across observers in **Table 3**. Meridian contributes the most variance.

### 82% of variance explained by Bouma law

Bouma (1970) discovered the linear relationship between crowding distance and eccentricity. He initially reported a slope of 0.5, which he later revised to 0.4 (Andriessen & Bouma, 1976). The Bouma law is:

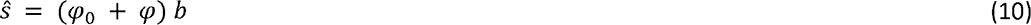

where s is crowding distance in deg, *φ* is radial eccentricity in deg, and *φ*_0_ (in deg) and *b* (dimensionless) are positive fitted constants (Bouma, 1970; Rosen et al., 2014). The dimensionless slope *b* is the Bouma factor. The horizontal intercept is –*φ*_0_, and the vertical intercept is *φ*_0_*b* (Liu & Arditi, 2000; Strasburger et al., 2011; Toet & Levi, 1992).

Crowding is one of several tasks whose threshold increases linearly with radial eccentricity, and such a task can be summarized by an *E*_2_ value that is the eccentricity at which threshold reaches twice its foveal value (Levi et al., 1985). In the Bouma law (**Eq. 10)**, *E*_2._ *φ*_0_.

Our large database of visual crowding thresholds (**Table 1**) is very well fit (*R*^2^ = 82.45%) by the two-parameter linear Bouma law (**Eq. 10**), showing that most of crowding’s variation in our data is explained by eccentricity. Just two degrees of freedom, *b* and *φ*_0_, suffice to fit all 650 data points (13 thresholds measured in each of 50 observers). The estimated slope*b* was 0.23, just over half of Bouma’s 0.4. Our database consists of measurements at five locations with radial and tangential flankers, and two fonts. To capture the effect of these parameters on the Bouma factor we propose an extended version of the Bouma law.

### 94% of variance is explained by extended Bouma law

Crowding depends on more than just eccentricity. Crowding varies substantially across meridians (right, left, up, or down), crowding orientation (radial or tangential), target kind (e.g., letters or symbols) and across individuals. Here, we enhance the Bouma law by including these other variables. One by one, the extensions add model parameters for meridian, crowding orientation, target kind, and observer. The models and the variance that they account for are summarized in **Table 4**.

*Meridian*. Factor *b*_0_, which allows *b* to depend on the meridian *θ* (right, left, up, or down)

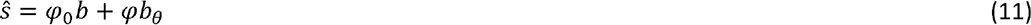

where *b* from **Eq. 10** now represents the*geometric mean* of *b*_0_, *b* = 10^(mean(log_10_(*b*_0_))). Note that the meridian is undefined at the fovea.

*Crowding orientation*. Factor *f*_dir_ depends on crowding orientation (radial or tangential).

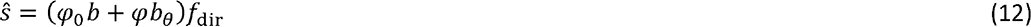

*Target kind*. Factor *t*_kind_ depends on target kind (e.g. Pelli or Sloan font).

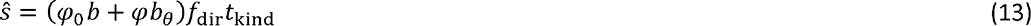

*Observer*. Finally, factor _<_ depends on the observer.

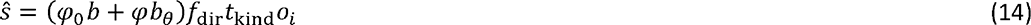

where ∏*_i_ o_i_* = 1. Adding factors to the original Bouma law accounts for more variance. Going from the simplest to the most enhanced model (**Eqs. 10 and 14**) increases explained variance from 82% to 94% (**Table 4**). Model performance is improved by adding the meridian factor (*R*^2^ = 89%) and crowding orientation (*R*^2^ = 93%). Adding the target-kind factor explains hardly any more variance, increased from 92.54% to 92.63%. Finally, the most enhanced model, with an observer factor, explains 94% of variance. The models are all cross validated, so the additional variance explained is not a necessary consequence of the increase in parameters. If the additional parameters were overfitting the training data, then we would find less variance accounted for in the left-out test data.

Since model parameters can be added in any order to form the final model (**Eq. 14**), we asked how much each parameter contributes to the total explained variance. For each parameter, we begin with the full model, remove that parameter, calculate the explained variance for the reduced model, and assess the drop in explained variance. We find that, after eccentricity, meridian contributes the most (4.5%) and target kind contributes the least (0.2%). (Note that most of our data were collected with one font; we expect target kind to explain more variance in studies that emphasize comparison of fonts, or other target kinds.) Results are shown in **Table 5**.

**Table 4.** How well the Bouma law and its extensions predict crowding distance. We begin with the Bouma law (Eq. 10). Successive models try to account for more variance by adding factors that depend on the meridian, crowding orientation, font, and observer. Each row gives a significantly better fit than the row above (assessed with F-test using Eq. 3). The R (Eq. 2) column shows cross-validated variance accounted for in predicting log crowding distance over the whole visual field (13 thresholds per observer). Pearson’s R shows the correlation between acquired and predicted data, and RMSE is root mean square error.

| Model | Equation | $R^2$ | $R$ | RMSE | No. of parameters |
| --- | --- | --- | --- | --- | --- |
| **Bouma law** | $s = (\varphi_0 + \varphi) b$<br>(10) | 82.45% | 0.90 | 4.80 | 2 |
| × **meridional factor** | $s = \varphi_0 b + \varphi b_\theta$<br>(11) | 88.96% | 0.94 | 3.80 | 5 |
| × **crowding orientation** | $s = (\varphi_0 b + \varphi b_\theta) f_{\text{dir}}$<br>(12) | 92.54% | 0.96 | 3.13 | 7 |
| × **target-kind factor** | $s = (\varphi_0 b + \varphi b_\theta) f_{\text{dir}} t_{\text{kind}}$<br>(13) | 92.63% | 0.96 | 3.11 | 9 |
| × **observer factor** | $s = (\varphi_0 b + \varphi b_\theta) f_{\text{dir}} t_{\text{kind}} o_i$<br>(14) | 93.86% | 0.97 | 2.84 | 59 |

**Table 5.**
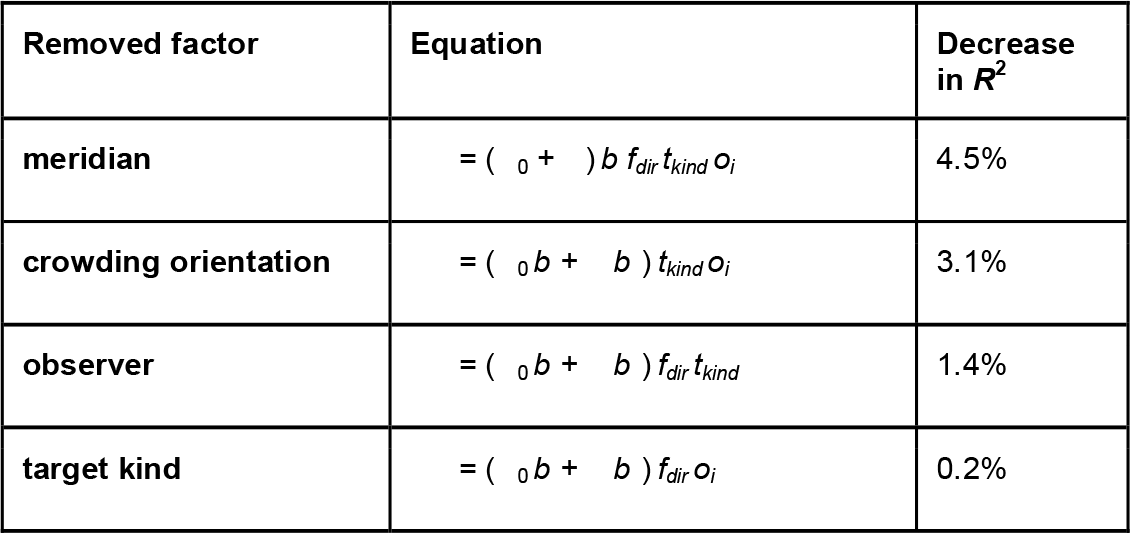
Parameter contribution to the most extended Bouma law. We measured the contribution of each parameter to the full model.

Since the enhanced model accounts for more variance than the original Bouma law, we looked in the data for systematic effects of these parameters. Figure 6 plots each set of model parameters after normalizing by the geometric mean of that set, where a set is the four meridians, two crowding orientations, two fonts, or fifty observers. Asymmetry within each factor is discussed in the next section. Except for target kind, each of the factors (meridian, crowding orientation, and observer) accounts for a roughly two-fold variation in the Bouma factor (dashed horizontal lines in **Fig. 6**).

### Radial Bouma factor varies twofold across meridians

Most of the thresholds in our dataset are for the Sloan font with radial flankers. Using these data, we explore the variation of Bouma factor across meridians. We estimate the Bouma factor by fitting **Eq. 10** for each participant and meridian independently (**Fig. 7A**). The Bouma factor is smallest along the right meridian and highest along the upper meridian (**Fig. 7B**). Bouma factor is 0.184 right, 0.237 left, 0.300 lower, and 0.381 upper meridian; overall geometric mean = 0.27).

*Meridional asymmetries.* We find three asymmetries. (As a reporting convention, we refer to the “advantage” of a smaller Bouma factor.) 1) Along the vertical midline, there is a 0.79 lower:upper advantage. 2) Along the horizontal midline, there is a 0.78 right:left advantage.. 3) Finally, there is a 0.62 horizontal:vertical advantage (based on geometric mean of the Bouma factors from right and left meridian vs upper and lower meridian). All reported asymmetries are highly consistent across participants (**Fig. 7C**). ANOVA reveals that there is a significant effect of meridian *F*(3,196) = 92.76, *p* < 0.001 and post-hoc analysis shows that Bouma estimates at each meridian are significantly different from each other (all *p* < 0.001; corrected for multiple comparisons). For each meridian we also estimate the eccentricity<_0_ at which the crowding distance reaches twice its foveal value. <_0_ was 0.37 deg ± 0.02 for right, 0.29 deg ± 0.02 for left, 0.22 deg ± 0.01 for lower and 0.17 deg ± 0.01 for upper meridian.

### Tangential Bouma factor is roughly half of radial

Unlike radial crowding, tangential crowding is the same in left and right meridians according to the Wilcoxon rank sum test (*z* = -0.73, *p* = 0.49). The standardized (see section on corrected Bouma factor below) tangential Bouma factor is much smaller than radial: 0.13 on the right and 0.14 on the left meridian. The tangential:radial ratio is 0.60 in the right meridian and 0.50 in the left.

### Bouma factor varies with target kind

Pelli and Tillman (2008) highlighted the remarkable degree to which crowding distance is conserved across stimulus kind, but later work shows that crowding distance does differ substantially between some target kinds (e.g. letters vs. symbols; Grainger et al., 2010). Along the horizontal midline, standardized Bouma factor for Sloan font was 0.239 on the right and 0.308 on the left, and slightly higher for the Pelli font: 0.325 on the right and 0.377 on the left. Overall, there is a 0.78 Sloan:Pelli ratio of standardized Bouma factors (geometric mean of the ratio taken at each meridian) and the difference between fonts was statistically significant (z = 3.58, p < 0.001). The model performance is slightly improved by adding the target-kind factor (**Eq. 13**). This factor contributed little to the overall variance explained by the model because most of the data came from trials with the same target kind (Sloan letters). So even though excluding target kind as a factor from the model caused inaccurate predictions for the Pelli font, the reduction in variance explained is negligible because nearly all of the dataset is based on one font. We anticipate that the target-kind factor will account for more variance in datasets that focus on comparing target kinds.

### Bouma factor varies twofold across observers

Bouma factor varies with meridian, crowding orientation, and target kind. Here, in this section, we quantify differences between observers. First, we estimated how well the Bouma law fits individual-participant data. Fitting **Eq. 10** to the right meridian data for each participant results, on average, in 97% explained variance, confirming that individual crowding data are well described by the linear model. Next, for each observer we fit the whole model to estimate the observer’s overall Bouma factor (**Fig. 8**). We also report individual differences in acuity. Individual differences are characterized by the standard deviation of the log of the threshold. Radial Bouma factor for the Sloan font varies approximately two-fold across observers (SD of log *b* = 0.08). This variation is unchanged for tangential flankers (SD of log *b* = 0.08) and nearly doubles for the Pelli font (SD of log *b* = 0.11). Foveal acuity *a* and foveal crowding distance *s* also vary two-fold. For crowding, the*φ*_0_ values also vary two-fold and range between 0.17 and 0.37 (Song et al., 2014). We also report the SD of the retest minus test difference for the Bouma factor estimated with radial flankers and Sloan font (**Fig. 8B**). For each observer we fit one Bouma factor for the test session and one Bouma factor for the retest. Differences across observers are much larger than those of test-retest. The 0.08 SD of the log Bouma factor across observers is 3 times larger than the 0.03 SD of test and retest, showing that one measurement is enough to distinguish individual differences.

### Supralinearity: Bouma factor increases with eccentricity

Bouma discovered the linear increase of crowding distance with eccentricity. We have seen that this linear equation fits our data well. However, seeing that we have 50 participants and data at 0 to 10 deg, a reviewer suggested that we examine how well the Bouma factor is conserved across eccentricity. To estimate the Bouma factor we fit **Eq. 10** for Sloan font with radial flankers to our data at 0 deg plus either 5 or 10 deg (**Fig. 9A**). On average Bouma factor is 1.4 higher at 10 than 5 deg eccentricity. This effect was statistically significant (*F*(1,398) = 42.3 *p* < 0.001), and there was no interaction between eccentricity and meridian (*F*(3,392) = 0.513 *p* = 0.674). This shows that the growth of crowding distance with eccentricity is actually more than linear. Indeed, the Coates et al. reanalysis of Bouma 1970 shows a similar supralinearity. Motivated by this finding, we invited 10 observers already in the main dataset (0, 5, 10 deg) to also measure crowding distance at 20 and 30 deg eccentricity.

When fitting data, there is a long tradition of using the shortest polynomial that fits adequately. Bouma (1970) initially suggested proportionality, with one degree of freedom. Measurements of nonzero crowding distance at 0 deg eccentricity led to a linear equation, with two degrees of freedom. Seeing curvature in our data from 10 observers from 0 to 30 deg, we enhanced the Bouma law from linear to quadratic (three degrees of freedom) to fit the data. **Eq. 15** adds a quadratic term to **Eq. 7** to allow the slope to grow with eccentricity. Replacing **Eq. 10** by **15** increases the degrees of freedom from 2 to 3, and increases the explained variance from 90% to 95% (**Fig. 9B-C**), 

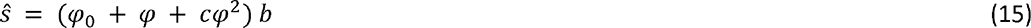

where ŝ is predicted crowding distance (in deg), < is radial eccentricity (in deg), and < (in deg), *b*, and *c* are degrees of freedom.

### Correlation of log crowding distance across visual field, crowding orientation, and target kind

We explored the pattern of correlations of log crowding distance for visual field locations, crowding orientations, and target kinds. These correlations are shown in **Figure 10A** where each cell shows Pearson’s *r* between two measurements. Rows are sorted so that the average correlation decreases from top to bottom. We find that log crowding distance measured on the right meridian at 10 deg eccentricity with Sloan font and radial flankers yields the highest average correlation with other log crowding distances (*r* = 0.39 with all, and *r* = 0.41 when fovea is excluded). Foveal log crowding distance measured with Pelli font yields the smallest average correlation with the rest of the log crowding distances.

To summarize how correlation depends on stimulus properties we estimated the average correlation across measurements when 1, 2 or 3 stimulus properties (eccentricity, meridian, target kind, crowding orientation) are modified (**Fig. 10B**). The test-retest correlation of 0.54 is plotted at zero changes. The average correlation drops to 0.30 with 1 change, to 0.25 with 2 changes, and to 0.18 with 3 changes. We also estimated what is the average correlation at the same stimulus location, and we only vary font and crowding orientation (right or left meridian at 5 deg). We find an average correlation (across 2 changes) of *r* = 0.54. On the other hand, when we change the location only and keep stimulus properties (e.g., radial flankers, Sloan font, 5 deg) we get a much lower correlation of *r* = 0.32. This indicates that when correlating crowding distances, location matters more than any other stimulus property.

### Correlation of Bouma factor ***b*** and intercept *φ*_0_

The Bouma law has two degrees of freedom, *φ*_0_ and *b*, which are anticorrelated, *r* = -0.51 (geometric mean across meridians).

### Standarized Bouma factor and its asymmetries across different studies

Visual field asymmetries can help identify the neural origin of perceptual phenomena (Afraz et al., 2010; Himmelberg et al., 2023). We compare our estimates of the Bouma factor to all the previous studies that measured crowding asymmetry.

*Estimating slope from just one point.* The Bouma law has two degrees of freedom, the slope *b* and the negative intercept <_0_. Estimating two parameters requires two measurements but many crowding studies report crowding distance at only one eccentricity. In the complete case, we have thresholds *S*_0_ and *S* at eccentricities 0 and *φ*, and we use the definition of the Bouma factor *b* as the slope*b* = (*s*- *s*_0_)/(*φ*- 0 . In the incomplete case, we have only threshold *S* at eccentricity *φ*. One might try to estimate the missing foveal threshold *s*_0_ or negative intercept *φ*_0_, but the simplest thing to do is to neglect *φ*_0_ (pretend it’s zero), and estimate *b*^ = s*⁄ φ*. The estimate has fractional erro ε = (*b*^ - *b*) *⁄ b* = *φ*_0_ *⁄ φ.* Thus neglecting *φ*_0_, possibly because the foveal threshold is unknown, leads to a fractional error *φ*_0_*⁄ φ*. The studies in Table 2 used eccentricities *φ* ≥ 2 deg. Our measurements estimate *φ* = 0.24 deg. Thus at 2 deg or beyond, the fractional error in estimated Bouma factor will be at most 0.24/2 = 12%. The fractional error drops to 5% at 5 deg and 2% at 10 deg.

Each Bouma factor was multiplied by a correction factor to account for criterion differences and log vs. linear flanker spacing (**Table 2**). Correction provides a *standardized* Bouma factor *b*’ for each study (**Table 6**). **Figure 11** compares crowding across studies by plotting the standardized Bouma factor vs. meridian.

**Figure 11.**
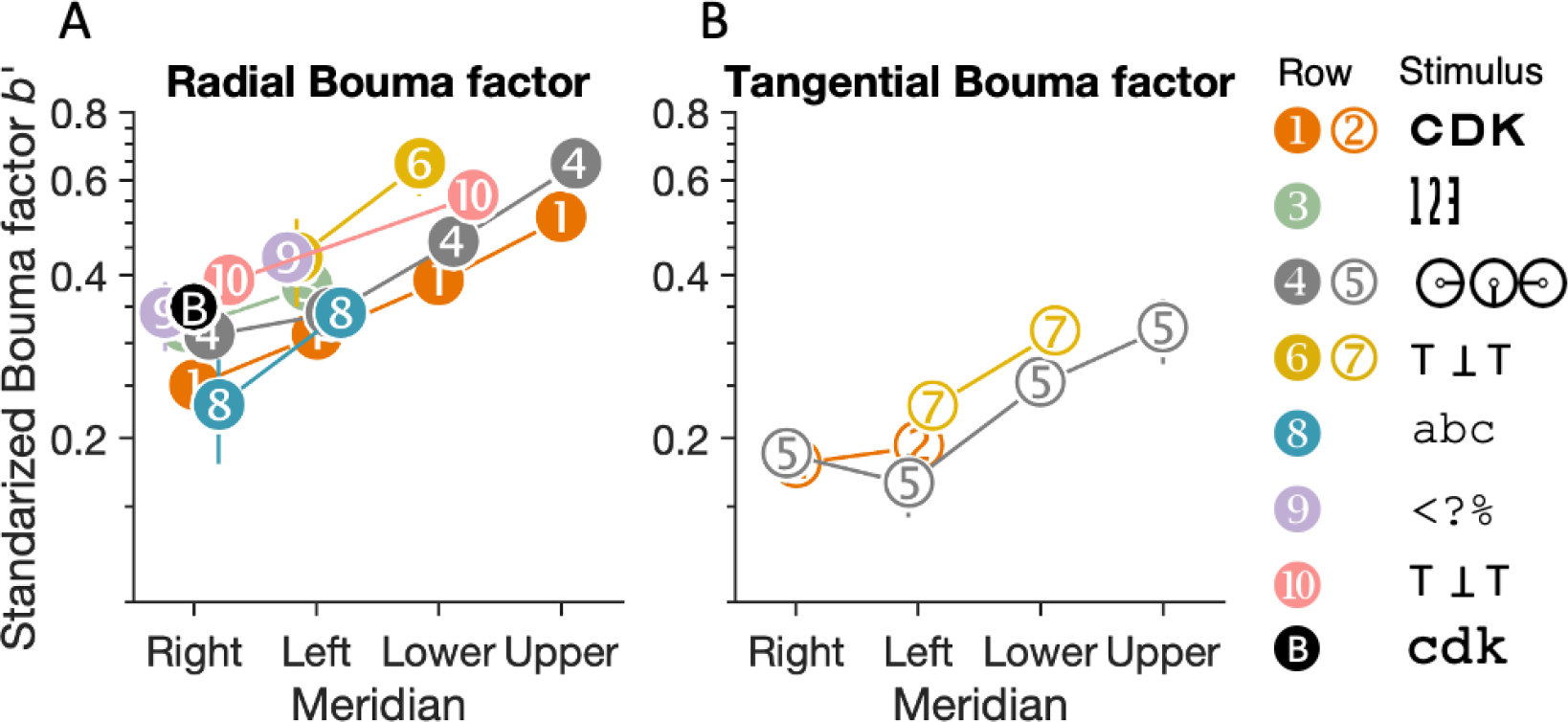
Standardized Bouma factor vs. meridian for various studies and target kinds. Each numbered point corresponds to a numbered row of data in Table 6. **A)** Comparison of radial-crowding studies and **B)** tangential-crowding studies. Both panels plot standardized Bouma factor vs. meridian. The legend shows the crowding stimuli. The white-on-black *B* symbol is the standardized Bouma factor estimated from Bouma (1970) data at 4 deg eccentricity with a 75% threshold criterion, with help from the reanalysis in Coates et al. (2021).

**Table 6.**
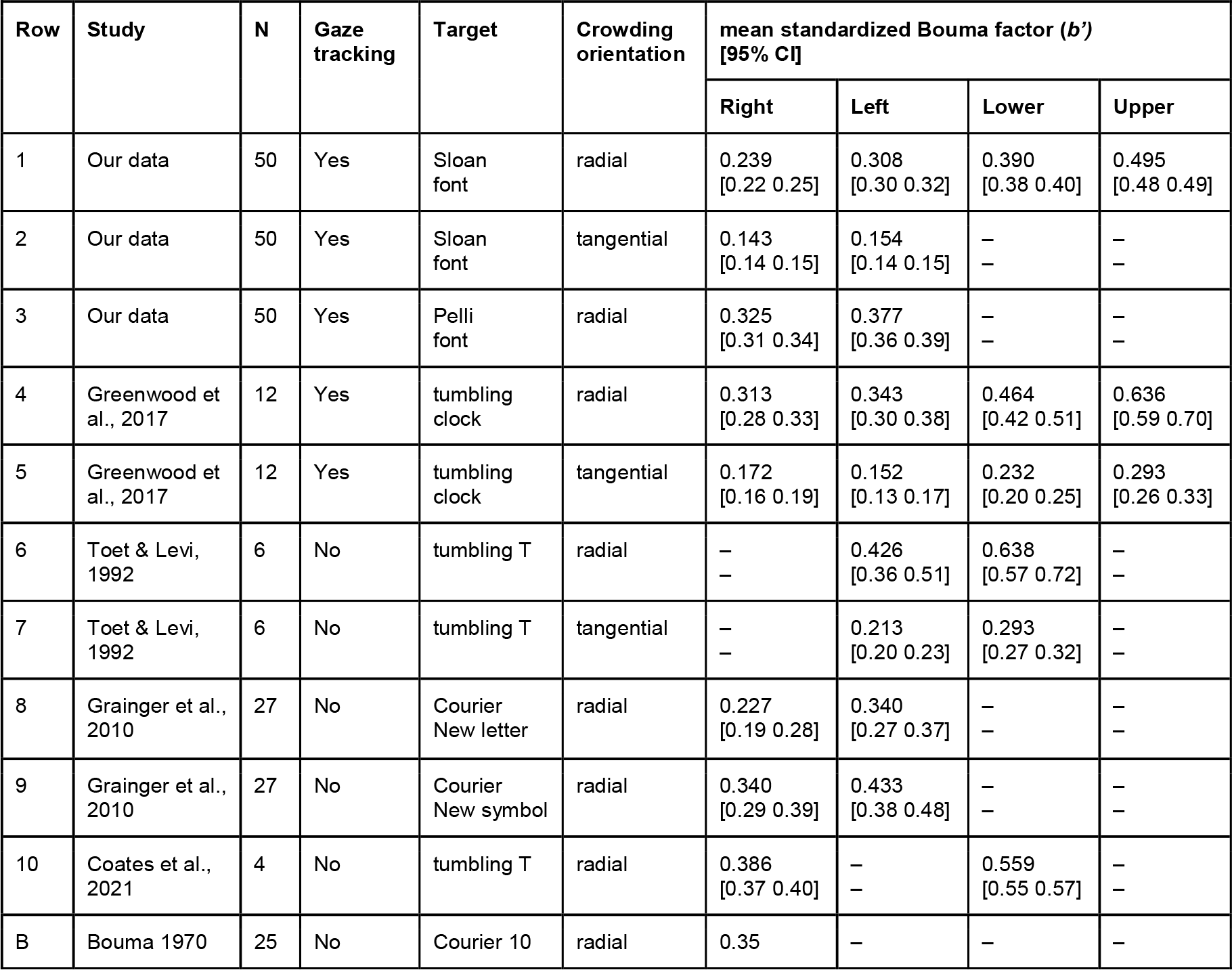
Standardized Bouma factor vs. meridian across studies. We compare our results with those of 4 studies that estimated crowding distance on at least two of the four cardinal meridians and report the Bouma factor by dividing crowding distance by the target’s eccentricity. We standardized each Bouma factor by correcting for differences in number of target choices and threshold criterion (see last column of Table 2). Data from Greenwood et al. (2017) and Coates et al. (2021) were obtained through personal communication. Data from Toet and Levi (1992) and Grainger et al. (2010) were digitized from figures in their papers. Data from Bouma 1970 were replotted from Coates et al. (2021). We estimated the standard error of Bouma factors in the Grainger et al. (2010) study based on the p-value they reported for meridian differences.

| Row | Study | N | Gaze tracking | Target | Crowding orientation | mean standardized Bouma factor ($b'$) [95% CI] | | | |
| --- | --- | --- | --- | --- | --- | --- | --- | --- | --- |
|  |  |  |  |  |  | Right | Left | Lower | Upper |
| 1 | Our data | 50 | Yes | Sloan font | radial | 0.239<br>[0.22 0.25] | 0.308<br>[0.30 0.32] | 0.390<br>[0.38 0.40] | 0.495<br>[0.48 0.49] |
| 2 | Our data | 50 | Yes | Sloan font | tangential | 0.143<br>[0.14 0.15] | 0.154<br>[0.14 0.15] | – | – |
| 3 | Our data | 50 | Yes | Pelli font | radial | 0.325<br>[0.31 0.34] | 0.377<br>[0.36 0.39] | – | – |
| 4 | Greenwood et al., 2017 | 12 | Yes | tumbling clock | radial | 0.313<br>[0.28 0.33] | 0.343<br>[0.30 0.38] | 0.464<br>[0.42 0.51] | 0.636<br>[0.59 0.70] |
| 5 | Greenwood et al., 2017 | 12 | Yes | tumbling clock | tangential | 0.172<br>[0.16 0.19] | 0.152<br>[0.13 0.17] | 0.232<br>[0.20 0.25] | 0.293<br>[0.26 0.33] |
| 6 | Toet & Levi, 1992 | 6 | No | tumbling T | radial | – | 0.426<br>[0.36 0.51] | 0.638<br>[0.57 0.72] | – |
| 7 | Toet & Levi, 1992 | 6 | No | tumbling T | tangential | – | 0.213<br>[0.20 0.23] | 0.293<br>[0.27 0.32] | – |
| 8 | Grainger et al., 2010 | 27 | No | Courier New letter | radial | 0.227<br>[0.19 0.28] | 0.340<br>[0.27 0.37] | – | – |
| 9 | Grainger et al., 2010 | 27 | No | Courier New symbol | radial | 0.340<br>[0.29 0.39] | 0.433<br>[0.38 0.48] | – | – |
| 10 | Coates et al., 2021 | 4 | No | tumbling T | radial | 0.386<br>[0.37 0.40] | – | 0.559<br>[0.55 0.57] | – |
| B | Bouma 1970 | 25 | No | Courier 10 | radial | 0.35 | – | – | – |

*Effects of meridian and target kind.* Two rows in **Table 6** — 1 (Sloan letter) and 4 (Tumbling clock) — report the radial standardized Bouma factor for all four cardinal meridians.**Figure 11A** shows that although the standardized Bouma factor is higher for the clock than for Sloan, by a factor of 1.3 (ratio of means for the right meridian 0.313/0.239 = 1.3), the two curves are otherwise similar, showing the same dependence on meridian. The 1.3:1 difference is not an artifact of number of choices or threshold criterion (**Table 2**). Both studies used gaze tracking to exclude fixation errors, so the difference is not a consequence of bad fixation. Thus, this seems to be a real 1.3:1 difference in standardized Bouma factor between target kinds, precisely what the target-kind factor *t*_kind_ is meant to account for in **Eq. 13**. The tumbling clocks may be more like each other than the 9 Sloan letters are and therefore produce larger crowding distance. Almost all other studies (Rows 3, 6, 9, 10) cluster above the Sloan font and show the same dependence on meridian. In general, we find that Courier New letters (Row 8) produce the smallest radial standardized Bouma factor (0.23 on the right meridian) and Tumbling Ts (Row 10) produce the largest radial standardized Bouma factor (0.39 on the right meridian).

*Tangential crowding.* We also compared standardized Bouma factors estimated with tangential flankers across studies (**Fig. 11B**). The tangential Bouma factor did not vary as much as radial, especially in the left meridian (**Fig. 11B**, Rows 2,5,7). Radial crowding estimates, even with the same stimuli, showed more variation in the standardized Bouma factor (**Fig. 11A**, Rows 1, 4, 6). Although our data did not show any difference between right and left meridians (Row 2), data extracted from Greenwood et al. (2017) do show a slight right:left advantage (Row 5).

*Meridional asymmetries.* **Table 7** and **Figure 12** report three Bouma-factor asymmetries (horizontal:vertical, right:left, and, lower:upper Bouma-factor ratios) for our and four selected studies. On average, the Bouma factor asymmetry is larger radially than tangentially. Radially, there is an advantage of horizontal over vertical meridian, right over left meridian, and lower over upper meridian in every study. The horizontal:vertical advantage seems to be insensitive to object kind as the estimates are clustered around ratio of 0.6-0.7. Similarly, lower visual field advantage is close to 0.8 for both studies that tested at this location (Rows 1 and 4). The right:left asymmetry is the most variable. The right:left ratio is smallest for Courier New letters (Row 8) and largest for clocks (Row 4).

**Figure 12.**
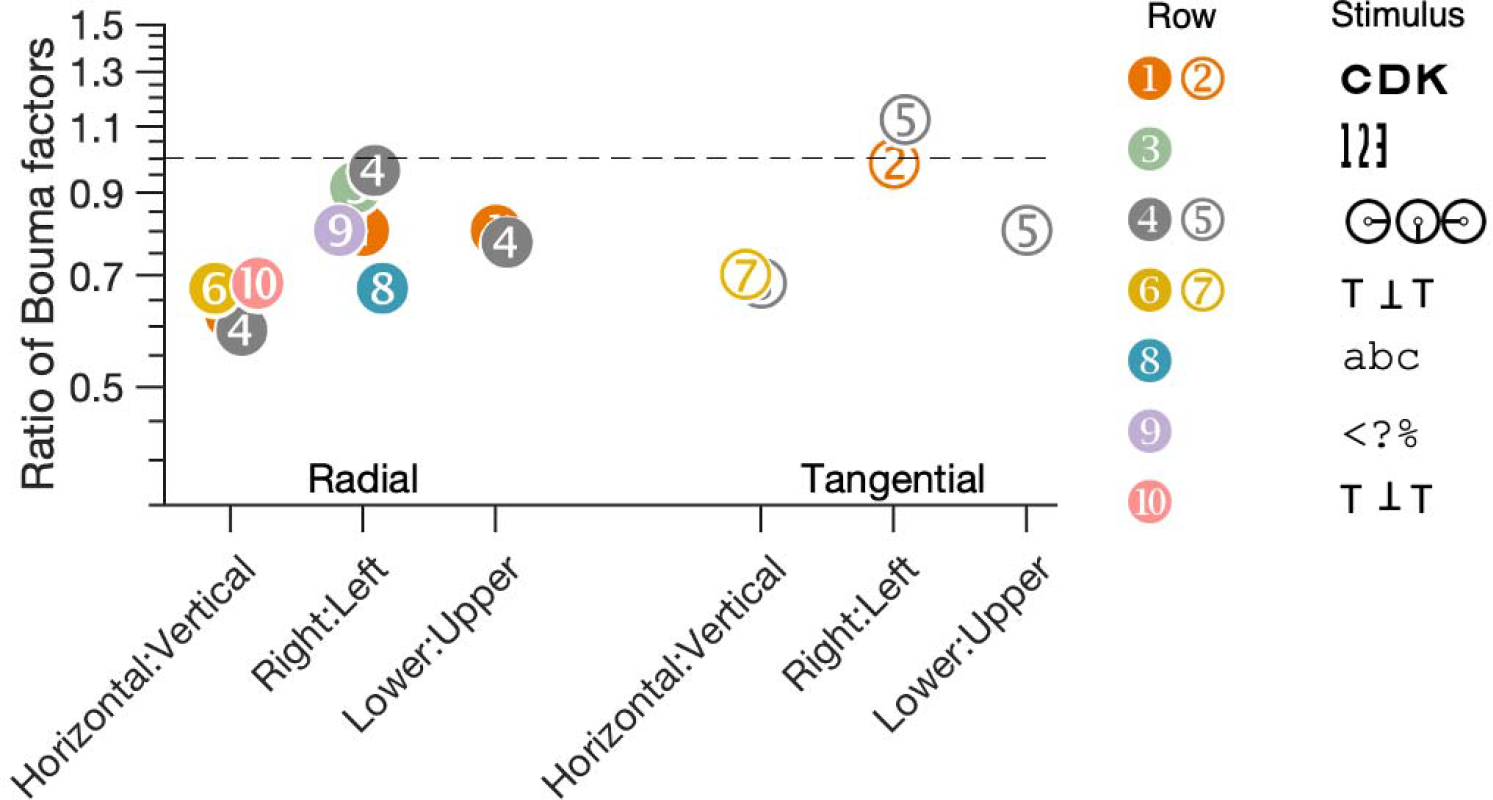
Plot of the three asymmetries, expressed as the ratio of Bouma factors. Each numbered point corresponds to a numbered row of data in Table 7. The horizontal dashed line at 1 represents no asymmetry. Each point is the ratio between Bouma factors. As in **Figure 11**, the legend shows examples of the objects that were used to measure crowding distance.

**Table 7.** Bouma factor and its asymmetries, compared with the literature. As in Table 6, we compare our results with those of studies that measured crowding distance on more than one of the four cardinal meridians. Each row lists the target type, crowding orientation, and mean ratio with standard-error intervals across participants.

| Row | Study | N | Gaze tracking | Target | Crowding orientation | Ratio of Bouma factors [95% CI] |  |  |
| --- | --- | --- | --- | --- | --- | --- | --- | --- |
|  |  |  |  |  |  | Horizontal:vertical | Right:left | Lower:upper |
| 1 | Our data | 50 | Yes | Sloan font | radial | 0.62 [0.60 0.64] | 0.78 [0.78 0.91] | 0.79 [0.76 0.82] |
| 2 | Our data | 50 | Yes | Sloan font | tangential | – | 0.93 [0.93 0.96] | – |
| 3 | Our data | 50 | Yes | Pelli font | radial | – | 0.86 [0.82 0.90] | – |
| 4 | Greenwood et al., 2017 | 12 | Yes | tumbling clock | radial | 0.59 [0.57 0.64] | 0.91 [0.88 1.05] | 0.73 [0.70 0.84] |
| 5 | Greenwood et al., 2017 | 12 | Yes | tumbling clock | tangential | 0.68 [0.6 0.75] | 1.13 [0.97 1.40] | 0.79 [0.73 0.91] |
| 6 | Toet & Levi, 1992 | 6 | – | tumbling T | radial | 0.67 [0.61 0.78] | – | – |
| 7 | Toet & Levi, 1992 | 6 | – | tumbling T | tangential | 0.73 [0.67 0.84] | – | – |
| 8 | Grainger et al., 2010 | 27 | – | Courier New letter | radial | – | 0.67 [0.56 0.76] | – |
| 9 | Grainger et al., 2010 | 27 | – | Courier New symbol | radial | – | 0.79 [0.70 0.90] | – |
| 10 | Coates et al., 2021 | 4 | – | tumbling T | radial | 0.69 [0.66 0.70] | – | – |

### What does peripheral crowding distance add to foveal acuity?

*Not predicted by acuity.* Any evaluation of the usefulness of crowding distance as a biomarker must assess what crowding tells us about the observer over and above what can be gleaned from foveal acuity, which is routinely measured in all optometric and ophthalmic exams. For our 50 observers, foveal acuity failed to predict peripheral crowding, with an insignificant average correlation of 0.04 (**Fig. 13;** grey peripheral circles). More generally both acuity and crowding measured in the fovea fail to predict peripheral crowding (average foveal-peripheral crowding correlation is an insignificant correlation of 0.15 – red circles).

**Figure 13.**
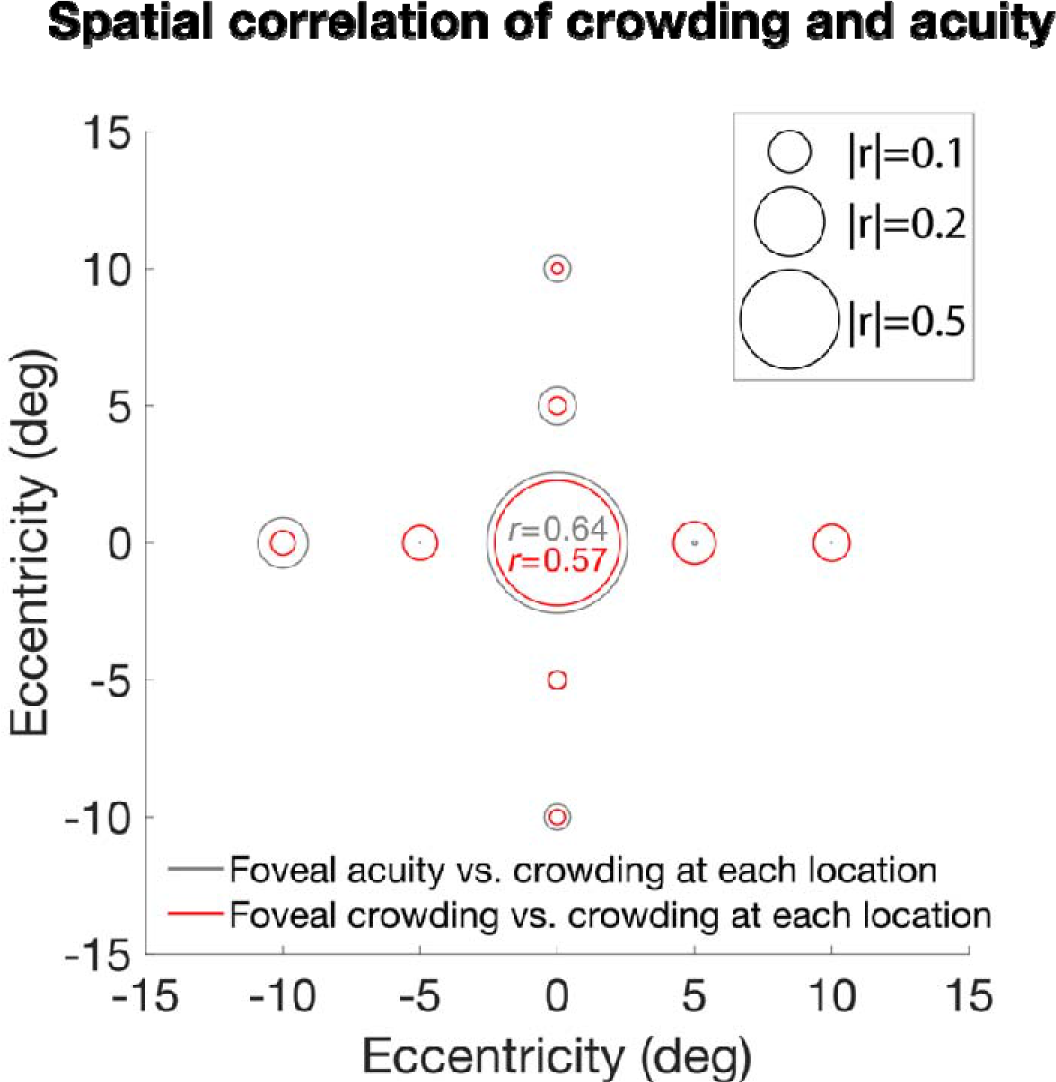
Foveal acuity and crowding fail to predict peripheral crowding. Correlation of foveal acuity (grey) and foveal crowding (red) with crowding everywhere. The central circles are test-retest for acuity (black) and crowding (red).

Within the fovea, we do find a significant correlation between acuity and crowding (*r* = 0.64). Thus, foveal acuity predicts foveal crowding but not peripheral crowding. If peripheral crowding is of interest, e.g., as a possible limit to reading speed, then it should be measured, since it is not predicted by foveal acuity.

*Foveal acuity and crowding.* Our procedure measures threshold by covarying size and spacing. Since we found that foveal acuity and crowding are correlated, one might ask whether the correlation is due to the measured crowding threshold being contaminated by acuity limits. For five observers, using the same CriticalSpacing.m software, Pelli et al. (2016) measured foveal spacing threshold with the Pelli font with several spacing:size ratios and, for each observer, confirmed that all the measured thresholds correspond to one spacing at different sizes. For those five normally-sighted observers, this showed that the procedure measured a crowding threshold, not acuity.

### The peeking-observer model

*Without gaze tracker.* We wondered how the Bouma factor estimate depends on fixation accuracy and we wondered if fixation accuracy might explain the difference between the two Bouma factor histograms in **Figure 1**. Peripheral identification is hard, so, in ordinary life, we typically first foveate a peripheral target that we need to identify. Despite instructing observers to fixate on the central cross, we know that gaze could be elsewhere during the target presentation. The observer is torn between the desire to follow the instruction to fixate the cross and the natural impulse to fixate an anticipated peripheral target location. When the target location is randomly one of several peripheral locations, the participant’s anticipation of location is often wrong. We model the participant’s gaze position by two distributions, one without and one with peeking. First, for the no-peeking *awaited-fixation* distribution, we used the measured eye position in the *awaited-fixation* dataset, in which the participant’s gaze was within 1.5 deg of the crosshair center for 250 ms immediately before target presentation, and we discarded trials in which gaze was more than 1.5 deg from the crosshair center during target presentation. The awaited-fixation gaze-position distribution is compact and roughly centered on the fixation crosshair. Second, we consider peeking toward a possible target location. We suppose that the participant peeks on a fraction *p* of the trials, and that the peeking eye movement travels only a fraction *k* of the distance from the crosshair to the possible target location, with a gaussian error (0.5 deg SD in x and in y). The *peeking distribution* has a mode corresponding to each possible target location, but at a fraction *k* of the possible target eccentricity. Gaze position is randomly sampled from the *peeking* distribution on a proportion *p* of trials and otherwise from the *awaited*-fixation distribution.

In the spirit of the Bouma law, our peeking-observer model assumes that the probability of identifying the target is given by a psychometric function

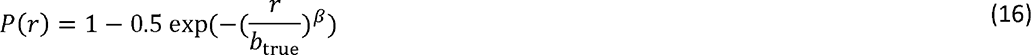

that depends solely on the ratio *r* of target-flanker spacing to actual target eccentricity, where *b*_true_ is the true Bouma factor and the steepness *β* is 2.3. For simplicity, the model omits threshold criterion and finger-error probability delta. Bouma factor *b* is estimated by 35 trials of Quest, assuming the true psychometric function, with a prior guess=0.11 of *r* and an assumed SD=2 of log *r*.

*Awaited-*fixation distribution. The *awaited-*fixation distribution was 3500 actual gaze positions at stimulus onset (35 trials x 50 participants x 2 sessions) measured with our EyeLink eye tracker in our *awaited-*fixation dataset. Recall that the stimulus was presented only once the gaze had been within 1.5 deg of the crosshair center for 250 ms.

*Peeking distribution*. We considered 1, 2, and 4 possible target locations (**Fig. 14A**). First, the target was always presented at one location (right meridian at 5 deg). Second, the target was randomly presented at ±5 deg on the horizontal midline. Third, the target was at 5 deg radial eccentricity on a random one of the four cardinal meridians (right, left, upper, lower). When participants “peek” a possible target location, depending on the number of possible locations, they have a 100%, 50%, or 25% chance of selecting the target location. Target and gaze position together define the actual target eccentricity.

**Figure 14.**
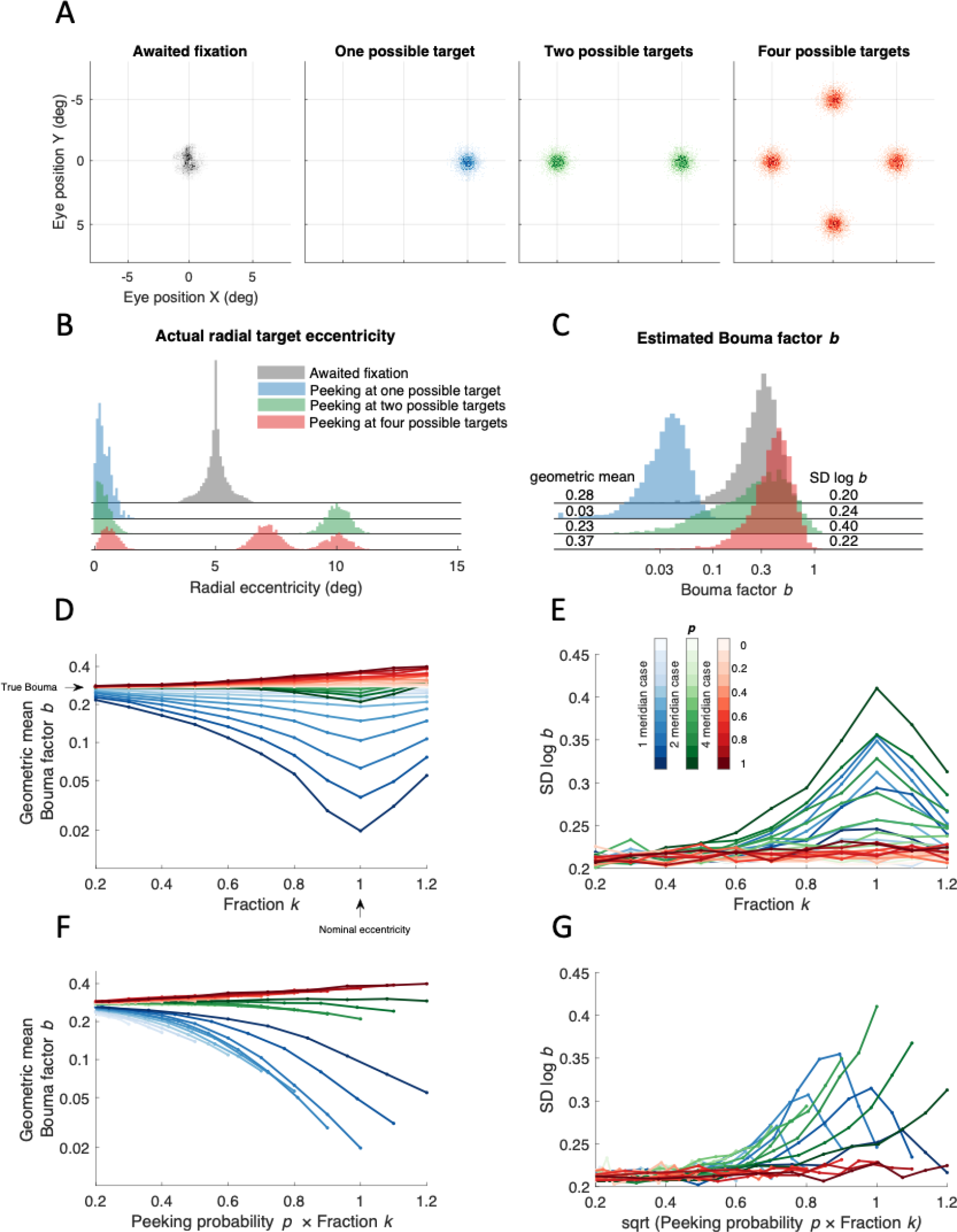
The peeking model. A) Scatter diagrams of the distribution of gaze position for each number of possible target locations. The gray histogram was measured by the EyeLink eye tracker in the *awaited-fixation* dataset. The green, blue, and red distributions of gaze position were synthesized assuming a full peek fraction *k*=1, and a gaussian (SD = 0.5 deg in x and 0.5 deg in y) centered at each possible target location **B)** Four histograms showing “actual” radial eccentricity of the target based on the distributions in panel A. (“actual” indicates that target eccentricity is computed relative to gaze position rather than the crosshair that the observer was asked to fixate.) **C)** 5000 Bouma factors estimated using Quest (35 trials) where each trial used the actual eccentricity. **D)** Geometric mean of the Bouma factor for each simulated case plotted vs. Fraction *k*. Arrow on the vertical axis indicates true value of the Bouma that we assumed (0.3) and arrow on the horizontal axis shows nominal eccentricity (5 deg). **E)** SD of Bouma factor vs. fraction *k*. Color saturation indicates the percentage of trials in which observers’ peek (see inset colorbar). **F-G)** Same as Panel D and E, however, the geometric mean of Bouma factor and its SD are plotted vs combination of peeking probability *p* with fraction *k*.

*Nominal* eccentricity of the target is relative to the crosshair.*Actual* eccentricity of the target is relative to gaze position when the target is presented. Peeking near the target position will reduce the actual eccentricity to practically zero, while peeking another target location could result in an actual eccentricity greater than nominal.**Figure 14B** shows the actual target eccentricities calculated based on the four distributions of gaze position in **Figure 14A**.

The nominal radial eccentricity of the target is always 5 deg, so we simulate an observer with a threshold spacing of 1.5 deg by using a psychometric function provided by Quest (beta = 2.30, delta = 0.01, gamma = 0.11). Thus, our model assumes a Bouma factor of 1.5/5 = 0.3. This is the parameter we try to estimate. The point of this exercise is to evaluate how various methods estimate the true Bouma factor. We simulate a block of 35 trials using Quest to estimate the 70%-correct spacing threshold from which we calculate the Bouma factor. We repeat this many times to get a histogram of estimated Bouma factor (**Fig. 14C**) for *awaited fixation* (gray) and *peeking* (colored) distributions.

The modelling shows that the geometric mean estimated Bouma factor*b* is lowest (0.03) for peeking with one possible location and highest with four (0.37), given a true Bouma factor *b* = 0.30. With no peeking (*p*=0), gaze position is from the *awaited-*fixation distribution, and the model estimate of Bouma factor is 0.28, very close to the true value of 0.30. The standard deviation of log Bouma factor *b* is highest (0.40) with two possible locations and lowest (0.22) with four.

The error (deviation from the assumed Bouma factor*b* = 0.3, see arrow on the vertical axis in **Fig. 14D**) in estimating the Bouma factor grows with the proportion *p* and fraction *k* of peeking (**Fig. 14D**). Observing that the error of estimated Bouma factor grows proportionally with *k* and that its SD grows proportionally with *k*^2^ (note the parabolic shape in **Fig. 14E**), we produced new figures (**Fig. 14F-G**) showing that geometric mean *b* is roughly linear with *p* × *k* and SD log *b* with Jp x k.

Given *p*, *k,* and the true Bouma factor, our peeking-observer model predicts the estimated Bouma factor *b*. We estimate *p* and *k* by fitting the model using maximum likelihood optimization. The higher the log-likelihood the better the fit. We wanted to know how often (*p*) and how far (*k*) the observer peeks and what is the estimated Bouma factor *b.* In **Figure 15A-B**, the scatter and breadth of the log-likelihood distribution result in a broad confidence interval for the product *p* × *k*. To estimate the error of the fit, we bootstrapped it by removing 25% of data at each iteration (n = 100). For*unmonitored* fixation, **Figure 15A**, bootstrapped parameters are consistent with high peeking (0.5 < *p* × *k* ≤1) and reject no peeking (*p* × *k* = 0). For *awaited* fixation, **Figure 15B**, bootstrapped parameters are consistent with low peeking (*p* × *k <* 0.5) and reject high peeking (*p* × *k* = 1). For each dataset, **Figure 15E-F** show that the human data are well matched by the simulated histogram. The two geometric means of *b* match, as do the standard deviations of log *b*. Our peeking model of eye position and crowding predicts the estimated Bouma factor in both cases, showing that the *unmonitored* fixation results are well fit by high peeking and the *awaited* fixation results are well fit by low peeking. One simple model of eye position and crowding fits all our data.

**Figure 15.**
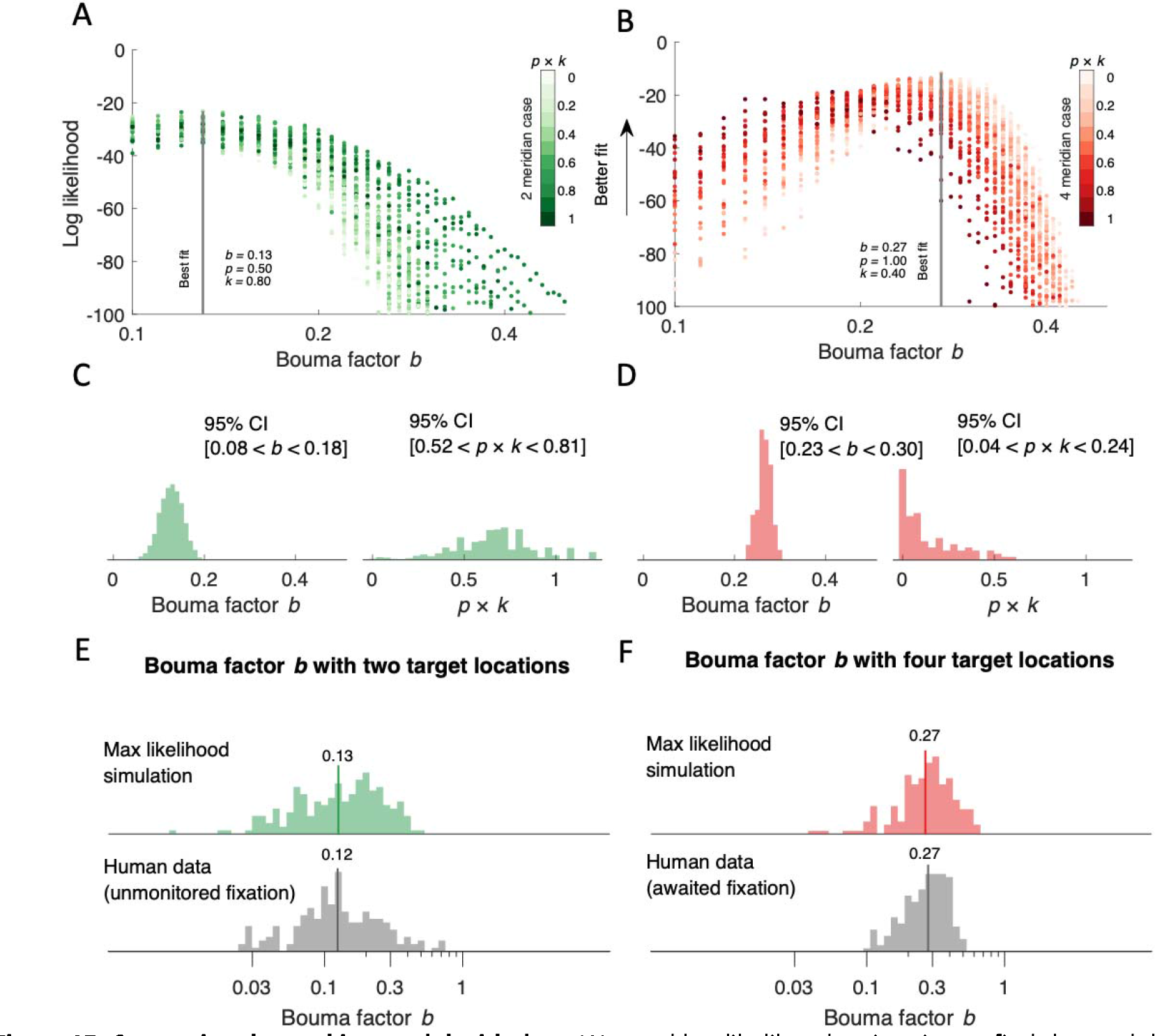
Comparing the peeking model with data. We used log-likelihood estimation to find the model parameter values that best fit our data. The higher the likelihood the better the fit. **A-B)** Log-likelihood is plotted vs. estimated Bouma factor *b*. Grey lines indicate best fit for which we plot model parameters. Brightness indicates the product *p × k*. **C-D**) Bootstrapped model estimates. For each model we bootstrapped the fit (n=100) by randomly removing 25% of data at each iteration and fitting the model on the remaining data. Each histogram contains 100 best fitted parameters from each iteration. The *p* × *k* confidence intervals were calculated after bootstrapping the *p* × *k* parameter (n=1000) and averaging 10 random samples at each iteration. This creates a normal distribution and allows the calculation of confidence intervals. **E-F)** Comparison of the histograms of best-fitting simulated and acquired data. The solid vertical line indicates the geometric mean. For *unmonitored* fixation, the geometric mean of Bouma factor *b* was 0.13 for the simulated data and 0.12 for the human data. The SD of log Bouma factor was 0.32 for simulated data and 0.31 for human data. For the *awaited* fixation geometric mean of Bouma factor was 0.27 for the simulated data and 0.27 for the human data. The SD of log Bouma factor was 0.21 for simulated data and 0.19 for human data. Note that the red distribution in panel D is different from the red distribution in **Fig. 1**. Here we plot data from all four meridians. Data plotted in **Fig. 1** are extracted only from the right and left meridians.

It is our impression that as observers gain experience with peripheral viewing, they peek less. Based on **Figure 15**, Bouma’s (1970) peeking rate must have been practically zero.

## DISCUSSION

Levi (2008), Pelli and Tillman (2008), Herzog et al. (2015), Strasburger (2020), and Coates et al. (2021) have reviewed the crowding literature. Most recently, Coates et al. (2021) provided a compact summary of the effects on the Bouma factor of contrast, size, target-flanker similarity and visual field location. This summary includes reanalysis of old data and shows a weak effect of stimulus duration. They also measured new data with two durations and two meridians confirming the effect of duration on the Bouma factor. Most of the data in their paper were acquired on fewer than 5 observers. We measured the effect of meridian, eccentricity, crowding orientation, and font with 50 observers. We did not measure effects of contrast, duration, or target-flanker similarity, but otherwise we confirm all the effects that they reported. We provide an equation predicting how crowding distance depends on meridian, target kind, and crowding orientation for each observer. We also show that crowding is reliable across days.

### The Bouma law and factor

*Bouma law*. The Bouma law describes the linear increase of crowding distance with eccentricity (Bouma, 1970, 1973; Levi, 2008; Pelli & Tillman, 2008; Rosen et al., 2014). The Bouma factor is the slope of that line (Rosen et al., 2014). The Bouma law is robust when fit to individual observer’s data (Pelli et al., 2004; Rosen et al., 2014; Strasburger, 2020; Strasburger et al., 1991). In this study, for the first time, we fit the Bouma law to data that include measurements from 50 observers tested with two crowding orientations at 9 locations of the visual field. The Bouma law is an excellent fit to our data and explains 82.5% of the variance despite being just a straight line with two degrees of freedom. We tried adding terms to the Bouma law to account for known factors: crowding orientation (Greenwood et al., 2017; Kwon et al., 2014; Petrov & Meleshkevich, 2011; Toet & Levi, 1992), meridional location of the stimulus (Fortenbaugh et al., 2015; Greenwood et al., 2017; He et al., 1996), target-kind (Coates et al., 2021; Grainger et al., 2010) and individual differences (Petrov & Meleshkevich, 2011; Veríssimo et al., 2021). We find that the enhanced model explains a bit more variance (increased from 82.5% to 94%). Eccentricity remains the dominant factor, accounting for 82.5% of the variance.

*Standardized Bouma factor*. We define the *standardized* Bouma factor *b*’ as the reported Bouma factor *b* (ratio of crowding distance to radial eccentricity) multiplied by a correction factor that account for differences in task from Bouma’s 25 choice alternatives, 75% threshold criterion, and linear flanker symmetry. Bouma reported a “roughly” 0.5 slope for radial letter crowding vs. eccentricity (Bouma, 1970). Andriessen and Bouma (1976) later reported a slope of 0.4 for crowding of lines. Coates et al. reanalyzed Bouma’s original data with various threshold criteria so we interpolated between the 70% and 80% thresholds to estimate the 75% threshold. Estimating the Bouma factor from Bouma’s original data using this criterion yields a Bouma factor of 0.35, in line with modern estimates of 0.3 (**Table 6** and Supplementary **Table 2**). **Figure 11A** shows that the corrected Bouma factor *b*’ ranges from 0.23 for Courier New letters to 0.39 for Tumbling T measured with radial flankers on the right meridian. That residual difference may be due to target kind (Coates et al., 2021; Grainger et al., 2010). This is further supported by our finding that the Bouma factor was 0.78 lower for the Sloan font than for the Pelli font.

### Supralinearity and the Bouma law

The linearity of the Bouma law implies that the Bouma factor is independent of eccentricity. The Coates et al. (2021) reanalysis of Bouma’s 1970 data found a twofold increase of the Bouma factor with eccentricity (from 1 to 7 deg), with a log-log slope of 0.35. Coates et al. speculated that this eccentricity dependence might be due to Bouma’s use of constant size stimuli at all eccentricities. Both acuity and crowding can limit measured thresholds for size and spacing across the visual field (Pelli et al., 2016; Song et al., 2014). If the threshold is independent of size, it is a crowding threshold and if the threshold is independent of spacing it is an acuity threshold. In 10 participants, we measured crowding distance at eccentricities of 0, 5, 10, 20, and 30 deg, scaling letter size with spacing as is now usual (see Methods). In our results with proportional letter size and controlled eye position, we find a similar twofold increase of the Bouma factor with eccentricity (from 5 to 30 deg), with a log-log slope of 0.38. Enhancing the Bouma law to allow a nonlinear dependence on eccentricity improves the fit to 10 observers’ data slightly, increasing the variance accounted for from 90% to 95%. This effect is small but detectable in data from 0, 5, and 10 deg (**Fig. 9A**) and becomes pronounced at eccentricities of 20 and 30 deg (**Fig. 9B**). To our knowledge, only a few past studies measured crowding beyond 10 deg eccentricity (Bouma, 1970; Kalpadakis-Smith et al., 2022; Kwon & Liu, 2019; Pelli et al., 2004) and all these datasets show supralinear growth with eccentricity. From the perspective of mathematical modeling Bouma initially suggested a simple proportionality with one term, which later was extended to linearity with two terms, and the evidence for supralinearity justifies a three-term quadratic polynomial. Biologically, it seems possible that the increase of the Bouma factor at high eccentricity reflects a compression of eccentric visual field in higher order areas.

Indeed, hV4 has a reduced peripheral representation when compared to earlier visual areas, V1, V2, and V3 (Arcaro et al., 2009; Goddard et al., 2011; Kolster et al., 2010; Winawer & Witthoft, 2015). This parallels the idea that the ventral visual stream, specialized in object recognition, emphasizes the central visual field (Levy et al., 2001; Ungerleider & Haxby, 1994).

### Crowding asymmetries

At any given eccentricity, the Bouma factor varies with polar angle. The Bouma factor is lower along the horizontal than vertical meridian (Greenwood et al., 2017; Petrov & Meleshkevich, 2011; Toet & Levi, 1992), is higher in the upper than lower meridian (Fortenbaugh et al., 2015; Greenwood et al., 2017; He et al., 1996; Toet & Levi, 1992), tends to be lower in the right than left meridian (Grainger et al., 2010; White et al., 2020) and approximately halves with tangential flankers (Greenwood et al., 2017; Kwon et al., 2014). In this work, we replicated all these asymmetries (**Fig. 12** and **Table 7**). The horizontal vs. vertical advantage and better performance in the lower vs. upper visual field is found for many visual tasks (Himmelberg et al., 2023), and these asymmetries parallel those found in population receptive field size, cortical magnification, retinal ganglion cell density, and the BOLD signal magnitude (Benson et al., 2020; Himmelberg et al., 2021; Kupers et al., 2022; Kupers et al., 2019; Kurzawski et al., 2022; Kwon & Liu, 2019; Liu et al., 2006; Silva et al., 2018). The right:left asymmetry seems to be least described and does not generalize across all tasks. Beyond crowding, right visual field advantages have been reported: For native readers of left-to-right written languages, like English, the right meridian outperforms left in word recognition (Mishkin & Gorgays, 1952). Worrall and Coles (1976) examined letter recognition across the visual field and found a significant right hemifield advantage only along the right horizontal midline. The similarities in asymmetry suggest a common mechanism, and the differences may be useful hints toward the cortical substrate of crowding.

### Standard deviation of measured acuity and crowding

To estimate the reliability of our measurements we acquired each threshold twice. Previous work showed improved performance in crowding tasks for repeated measurements (Chung, 2007; Malania et al., 2020). From their figures we estimate the second-block benefit to be 13% for Malania et al. and 20% for Chung. [Chung shows thresholds before and after 60 100-trial blocks. She shows percent correct for each block. By eye, we estimate that the benefit from first to second block is about a third that provided by the 60 blocks of training. Thus her 62% advantage (see average data from Table 1 – Chung 2007) after 60 blocks corresponds to the 20% advantage after the first block.] We find a modest second-threshold improvement for crowding thresholds measured with Sloan font at all tested locations and with the Pelli font in the fovea (less than 10%). Thresholds measured with Pelli font in the periphery yielded the highest improvement (23%). In our data the improvement is likely not due to acquiring familiarity with the task, since all observers participated in a training session, which consisted of repeated trials until 10 answers are correct. We find no improvement in acuity.

Overall, we find very good reproducibility of crowding and acuity thresholds **F**(**ig. 5**). The standard deviation of log Bouma factor *b* measured with Sloan font and radial flankers for test-retest is much lower than the standard deviation of log Bouma factor across observers (0.03 vs 0.08).

### Individual differences

Estimating individual differences requires data from many observers. In this paper we measured crowding in 50 observers, which is the biggest dataset of crowding measurements to date. Previous in-person crowding surveys included at most 27 observers (Grainger et al., 2010; Greenwood et al., 2017; Petrov & Meleshkevich, 2011; Toet & Levi, 1992). [An online crowding study tested 793 observers, but did not report individual differences (Liu et al., 2009).] To capture individual differences in the Bouma factor we included an observer factor *o*_i_ in the enhanced model. Adding the observer factor improved the explained variance from 92.6% to 94% (**Table 4**). Although this effect may seem negligible at first, we find that the Bouma factor varies twofold across observers, ranging from 0.20 to 0.38 (**Fig. 8A**). A similar, two-fold variation is observed for all other thresholds that we estimated (**Fig. 8A**).

*Bouma factor as a biomarker*. Large individual differences enhance crowding’s potential as a biomarker for studying cortical health and development. Specifically, crowding varies across children too (Kalpadakis-Smith et al., 2022) and predicts RSVP reading speed (Pelli et al., 2007). Foveal crowding distance drops threefold from age 3 to 8 (Waugh et al., 2018). If crowding correlates with reading speed of beginning readers, then preliterate measures of crowding might help identify the children who need extra help before they learn to read.

Measuring crowding distance across individuals in several diverse populations might expose any limit that crowding imposes on reading, yielding a norm for the development of crowding. Huge public interventions seek to help dyslexic children read faster and to identify them sooner. A virtue of crowding distance as a potential biomarker for dyslexia and cortical health is that it can be measured in 3.5 minutes.

### Crowding correlations

This paper reports 13 crowding thresholds for each of 50 observers. Such a comprehensive dataset allows for a correlation analysis to assess how well each crowding threshold predicts the others. We find a moderate correlation of crowding between peripheral locations (*r* = 0.39 averaged across all peripheral locations) and hardly any between fovea and periphery (*r* = 0.11). We also find that crowding measured with radial flankers correlates highly with crowding measured with tangential flankers at the same location (*r* = 0.53 for the right meridian, *r* = 0.50 for the left meridian). The threshold measurement that best predicts all other peripheral thresholds (excluding the fovea), with a correlation *r* = 0.41, is radial Sloan crowding at 10 deg in the right meridian.

*Effect of stimulus configuration vs. location*. We find higher correlation (*r* = 0.54) when the location is the same and the stimulus configuration is changed, than (r = 0.32) when stimulus configuration is the same and location is changed. Correlation of crowding distance depends more on location than configuration. Paralleling our result, Poggel and Strasburger (2004) found only a weak correlation across meridians for visual reaction times. Surprisingly little is known about spatial correlation of basic measures like acuity and contrast sensitivity.

### The peeking-observer model

We always asked the observer to fixate on the crosshair during each trial. We acquired data with two methods: *unmonitored fixation*, without gaze tracking, and *awaited fixation,* in which the stimulus was only presented when gaze was near the fixation cross. Both methods are described in detail in the Methods section. The two methods yield different Bouma factor distributions. Upgrading from *unmonitored* to *awaited* fixation increased the Bouma factor mean *b* from 0.12 to 0.20 and nearly halved the standard deviation of log *b* from 0.31 to 0.18. Histograms are shown in **Figure 1**. This peeking-observer model assumes firstly that performance on each trial depends solely on target eccentricity (relative to gaze position), secondly that the observer peeks on a fraction*p* of the trials and fixates near the crosshair on the rest of the trials, and thirdly that the location of the peek is a fraction*k* of the distance from the fixation mark to the anticipated target location.

In *unmonitored* fixation, the observer peeks with probability*p*. In *awaited* fixation, peeking is prevented by using gaze-contingent display and discarding any trials where gaze left the fixation cross while the target was present. Suppose there are two possible target locations. The Bouma factor distribution is unimodal for low values of *p* and becomes bimodal for high values for *p*. Our *unmonitored b* histogram is bimodal is best fit with a peeking probability of 50%. *Our awaited* fixation b histogram is unimodal and best fit by peeking restricted to the 1.5 deg from the crosshair allowed by the gaze tracker. Upgrading from the bimodal to the unimodal *b* distribution raised the mean *b* from 0.12 to 0.27 and nearly halved the standard deviation of log *b* from 0.31 to 0.19.

The peeking model does not account for the reduction of a target’s crowding distance that occurs in anticipation of a saccade to the target (Harrison et al., 2013). It is conceivable that on some *awaited*-fixation trials the observer was planning an eye movement to the correct target location and that this reduced crowding before the eye moved.

*Effects of duration and peeking*. Coates et al. (2021) reanalyzed crowding data from 16 studies and presented a scatter diagram of Bouma factor versus stimulus duration. The plot of Bouma factor vs. log stimulus duration had a semi-log slope of -0.16 describing how the Bouma factor drops with duration. Their analysis included many studies, with various threshold criteria, from various meridians, which introduced differences in the Bouma factor. To avoid these confounds, Coates et al. collected new data using a consistent threshold criterion and consistent locations. In their new results, increasing the duration from 67 to 500 ms decreased the Bouma factor by a factor of 1/1.6. However, none of these studies monitored fixation. Our **Figure 1** shows that, relative to controlled fixation, peeking can reduce the Bouma factor by a factor of 1.6 which is the size of the decrease with duration reported by Coates et al. If the probability of peeking grows with duration, then peeking might explain their drop in Bouma factor with duration.

### Why measure crowding?

*Peripheral crowding provides additional information about visual health*. Acuity is the threshold size of a target for recognition, while crowding is a spacing threshold. Clinical assessment routinely includes foveal acuity and not crowding. Both limit recognition of everyday objects. Our results show that peripheral crowding is independent of foveal acuity and might be a useful biomarker of visual health. Specifically, peripheral crowding might predict dyslexia (Bouma & Legein, 1977; Martelli et al., 2009; O’Brien et al., 2005). There are hints that crowding tends to be worse in dyslexia (Pelli et al., 2007). If crowding correlates with reading speed of beginning readers, then preliterate measures of peripheral crowding might help identify the children who need extra help before they learn to read.

*What about foveal crowding?* In healthy individuals, foveal crowding correlates with foveal acuity but there are some conditions in which the two are dissociated. Strabismic amblyopia makes crowding worse in the fovea, but not in the periphery (Song et al., 2014). This suggests that the fovea might be the most sensitive place to detect the increase in crowding associated with amblyopia. Traditional tests for crowding are mostly peripheral and use a fixation mark and a brief peripheral target, which are poorly suited for testing children and dementia patients whose attention may wander. Such participants will fixate much more reliably on a foveal target.

We hope there will be clinical studies to assess the diagnostic benefit of measuring crowding, which takes 3.5 minutes.

## CONCLUSIONS

1. The well-known Bouma law — crowding distance depends linearly on radial eccentricity — explains 82% of the variance of log crowding distance, cross-validated. Our enhanced Bouma law, with factors for observer, meridian, and target kind, explains 94% of the variance, cross-validated. The very good fit states the central accomplishment of the paper and shows how well the linear Bouma law fits human data.
2. The Bouma factor varies twofold across meridians, and radial vs. tangential crowding orientations.
3. Consistent with past reports, five asymmetries each confer an advantage expressed as a ratio of Bouma factors: 0.62 horizontal:vertical, 0.79 lower:upper, 0.78 right:left, 0.55 tangential:radial, and 0.78 Sloan font:Pelli font.
4. The Bouma factor varies twofold across observers. Differences across observers are much larger than those of test-retest. The 0.08 SD of log Bouma factor across observers is triple the 0.03 SD of test-retest, so one 3.5-minute threshold is enough to capture individual differences.
5. The growth of crowding distance with eccentricity is supralinear, but a linear fit is nearly as good, unless the range of eccentricities is huge.
6. Crowding distance measured at 10 deg eccentricity along the right meridian is the best predictor of average crowding distance elsewhere (*r* = 0.39).
7. Peripheral crowding is independent of foveal crowding and foveal acuity.
8. Peeking can be avoided by use of a gaze-contingent display. Peeking nearly halves the geometric mean Bouma factor *b*, and nearly doubles the standard deviation of log *b,* from 0.18 to 0.31.

## ACKNOWLEDGEMENTS

We thank reviewer Hans Strasburger for many helpful suggestions which led to significant improvements, including the standardized Bouma factor. Thanks to Nina Hanning for helping with EyeLink hardware and software and for providing code to extract eye position at each trial. Thanks to Larry Maloney for suggesting bootstrap analysis of log-likelihood. Thanks to Barbara Finlay for helpful discussion of how psychophysical crowding correlations might relate to retinal developmental. Thanks to Maria Pombo for ANOVA help. Thanks to Ashley Feng and Laura Suciu for comments on the manuscript. The authors contributed as follows: Jan Kurzawski collected data with *awaited* fixation, analyzed data with *awaited* and *unmonitored* fixation and wrote the paper; Augustin Burchell performed initial analysis of *unmonitored fixation* data. Darshan Thapa supervised testing for data with *unmonitored fixation*; Najib Majaj discussed, designed, analyzed, and wrote; Jonathan Winawer discussed, and helped design, analyze, and write; Denis Pelli directed the project, including design, software, data collection, analysis, and writing. Jan, Najib, and Denis met most days for a year to do the revisions.

## SUPPLEMENTARY MATERIALS

### Effect of equating either the linear or log spacing of two radial flankers

When measuring radial crowding, the target lies between two flankers on a radial line from fixation. Bouma spaced the flankers equally, and most investigators have followed suit.

However, we spaced the flankers symmetrically on a logarithmic rather than linear scale. This raises the question of how to compare crowding distances between experiments that spaced the flankers linearly vs logarithmically. Given the Bouma law (**Eq. 10**) and assuming that crowding distance depends primarily on the flanker-to-flanker distance, and negligibly on the target position between them, we show here that the crowding distance is expected to be 1.18 times larger when measured with linearly-spaced flankers than with log-spaced flankers.

Specifically, for a target at *φ* we choose the two flanker eccentricities *φ*_in_ and *φ*_out_ so that *φ* is located between *φ*_in_ and *φ*_out._

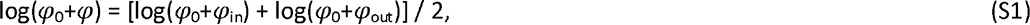

where *φ*_0_=0.15 deg, and we report the inner spacing *s* = *φ*-*φ*_in_. We can rearrange

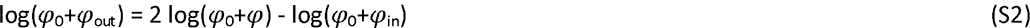

Thus, the two flankers are at different (linear) distances from the target, but we suppose that they are equally effective in crowding the target. We report the center-to-center spacing from the inner flanker to the target as the “spacing” *s*.

We now estimate the relation of crowding distances measured in these two ways. We suppose that degree of crowding is determined by the separation between flankers on the log scale, as expected from the Bouma law:

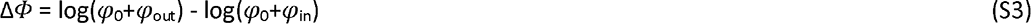

If we are in the periphery, i.e. *φ*_in_ > 1, then *φ*_0_=0.15 is negligible, and we can simplify,

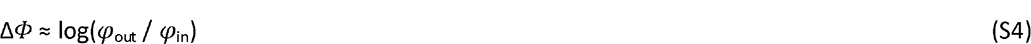

Of course, increasing spacing alleviates crowding, so the degree of crowding will drop as log spacing Δ*Φ* grows. Note that this model is at best an approximation, as it neglects position of the target. We are using it solely to compare crowding for two different ways of centering the target between flankers so the two target positions won’t differ by much.

*Linear flanker spacing*: With flankers spaced symmetrically about the target on a linear scale, both at distance *s* from the target:

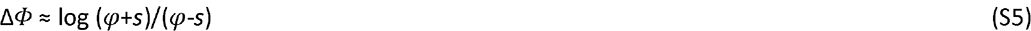

*Log flanker spacing*: With flankers spaced symmetrically on a log scale, the log flanker-to- flanker spacing is twice the log target-to-flanker spacing:

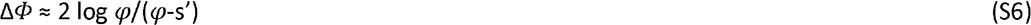

Now we equate the two log flanker spacings, one with linearly symmetric spacing*s*, the other with log-symmetric spacing *s’*.

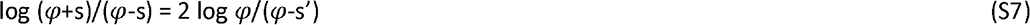

Solve for *s’*,

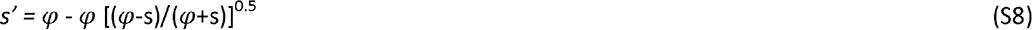

Now substitute *b*=*s*/*φ* and *b’*=*s’*/*φ,*

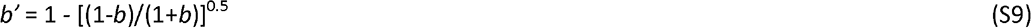

Figure S1 shows that, to a good approximation, this is a proportionality, with error of at most 0.014 over the relevant range 0 ≤ *b* ≤ 0.9,

**Figure S1.**
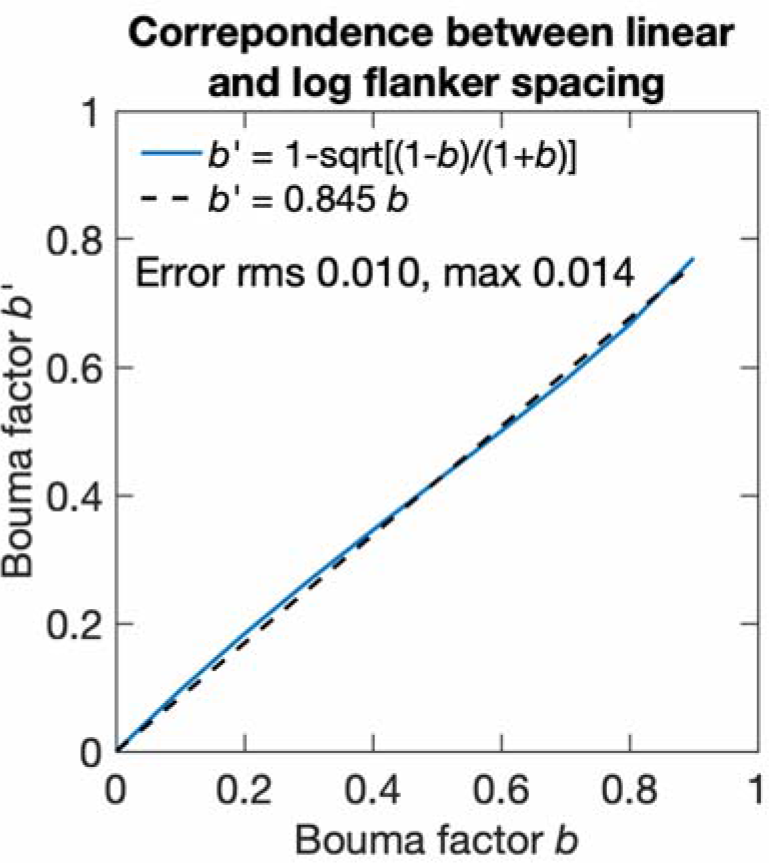
Log-symmetric spacing *s’*=*φb’* for same crowding effect as each linearly symmetric spacing *s* = *φb*.

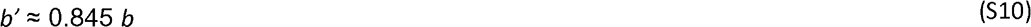

Thus, our log-symmetric spacing *s’=φb’* is approximately 0.845 times the linearly-symmetric spacing *s=φb* that is traditionally reported.

### Log-log versions of the Bouma models

The negative intercept *φ*0 is small and negligible at large eccentricity. (Zeroing it in the Bouma law produces less than 5% error in predicted crowding distance at eccentricities beyond 4.8 deg.) If we consider only peripheral results (>4.8 deg eccentric) we can set *φ*0=0, and express the Bouma models in log coordinates (Table S2). The multiplicative combination rule becomes additive in the new coordinates. As with the linear models, the cross-validated variance explained *R*^2^ increases with more parameters. At large eccentricity, these models are equivalent to the linear models presented in the main text, but fitting is quicker because the fitting error can be minimized by linear regression.

**Table S1.** Fitting the log-log version of the Bouma law (setting *φ*_0_=0 and modeling only peripheral data 2 *φ*>4.8 deg). Uppercase variables are the log10 of corresponding lowercase variables. R^2^ represents variance explained after model cross-validation. All our fitting minimizes error in log crowding distance so fitting the log-log version can be fit using linear regression.

| Model | Equation | $R^2$ (%) | Pearson's $R$ | No. of parameters |
| --- | --- | --- | --- | --- |
| Bouma law | $S \approx \phi + B$ | 53.53 | 0.78 | 1 |
| × meridional factor | $S \approx \phi + B_\theta$ | 71.38 | 0.85 | 4 |
| × crowding orientation | $S \approx \phi + B_\theta + F_d$ | 80.47 | 0.89 | 6 |
| × target-kind factor | $S \approx \phi + B_\theta + F_d + T_{kind}$ | 81.13 | 0.90 | 8 |
| × observer factor | $S \approx \phi + B_\theta + F_d + T_{kind} + O_i$ | 84.32 | 0.92 | 58 |

### Correcting the Bouma factor

Comparison of crowding measured with different threshold criterion, number of choices, and log vs. linear spacing of flankers is facilitated by calculating the Standardized Bouma factor *b*’, which corrects for these factors. Each row number in Table S2 corresponds to a row in Table 6. The correction factors come from Table 2.

**Table S2.** Radial Bouma factor before and after the correction. Row numbers correspond to numbers in Table 6. Each cell contains two numbers. The upper number is the Bouma factor before accounting for measurement differences and the lower one is the Standardized Bouma factor (already multiplied by the correction factor). Correction factors are calculated in Table 2. The Standardized Bouma factor is overall higher (e.g., 0.26 to 0.32 on the right meridian).

| Row | Correction factor | Bouma factor<br>Standardized Bouma factor |  |  |  |
| --- | --- | --- | --- | --- | --- |
|  |  | Right | Left | Lower | Upper |
| 1 | $\times 1.30$<br>(1.10×1.18) | 0.184<br>**0.239** | 0.237<br>**0.308** | 0.300<br>**0.390** | 0.381<br>**0.495** |
| 3 | $\times 1.30$<br>(1.10×1.18) | 0.25<br>**0.325** | 0.29<br>**0.377** | | |
| 4 | $\times 1.01$ | 0.31<br>**0.313** | 0.34<br>**0.343** | 0.46<br>**0.464** | 0.63<br>**0.636** |
| 6 | $\times 1.33$ | | 0.32<br>**0.426** | 0.48<br>**0.638** | |
| 8 | $\times 1.03$ | 0.22<br>**0.227** | 0.33<br>**0.340** | | |
| 9 | $\times 1.03$ | 0.33<br>**0.340** | 0.42<br>**0.433** | | |
| 10 | $\times 1.33$ | 0.29<br>**0.386** | | 0.42<br>**0.559** | |
| Geometric mean |  | 0.26<br>**0.30** | 0.32<br>**0.37** | 0.40<br>**0.50** | 0.49<br>**0.56** |

